# ARHGAP17 regulates the spatiotemporal activity of Cdc42 at invadopodia

**DOI:** 10.1101/2022.06.22.497207

**Authors:** Gabriel Kreider-Letterman, Abel Castillo, Eike K. Mahlandt, Joachim Goedhart, Agustin Rabino, Silvia Goicoechea, Rafael Garcia-Mata

## Abstract

Invadopodia formation is regulated by Rho GTPases. However, the molecular mechanisms that control Rho GTPase signaling at invadopodia remain poorly understood. Here, we have identified ARHGAP17, a Cdc42-specific RhoGAP, as a key regulator of invadopodia in breast cancer cells and by RhoGAPs characterized a novel ARHGAP17-mediated signaling pathway that controls the spatiotemporal activity of Cdc42 during invadopodia turnover. Our results show that during invadopodia assembly, ARHGAP17 localizes to the invadopodia ring and restricts the activity of Cdc42 to the invadopodia core, where it promotes invadopodia growth. Invadopodia disassembly starts when ARHGAP17 translocates from the invadopodia ring to the core, in a process that is mediated by its interaction with the Cdc42 effector CIP4. Once at the core, ARHGAP17 inactivates Cdc42 to promote invadopodia disassembly. Our results in invadopodia provide new insights on the coordinated transition between the activation and inactivation of Rho GTPases.

## Introduction

The risks associated with cancer metastasis makes breast cancer one of the leading causes of cancer related mortality among women. One way that metastatic cancer cells invade tissues is through the formation of protrusive, actin-rich structures called invadopodia, which mediate the degradation of the extracellular matrix (ECM) (Murphy and Courtneidge, 2011). The ability of cancer cells to form invadopodia has been correlated with their metastatic potential, and the loss of invadopodia with decreased tumorigenicity and dissemination (Meirson et al., 2018; Stoletov and Lewis, 2015; Yamaguchi, 2012). In addition, studies using intravital microscopy have documented the existence of invadopodia and confirmed their importance for intra/extravasation and dissemination in vivo (Gligorijevic et al., 2014).

The structure of invadopodia is typically comprised of an actin-rich core that is surrounded by a ring of adhesion and scaffolding proteins, like that of podosomes which occur in normal cells (Murphy and Courtneidge, 2011). Invadopodia form through a series of steps that is initiated by the disassembly of focal adhesions and stress fibers, and the recycling of many of their components into new invadopodia (Hoshino et al., 2012; Oikawa et al., 2008). Assembly of new invadopodia starts with the formation of actin- and cortactin-rich puncta, closely followed by the recruitment of adhesion proteins such as vinculin and paxillin (Hoshino et al., 2013). Invadopodia reach maturity upon the recruitment of matrix metalloproteases which facilitate matrix degradation (Hoshino et al., 2013). There is less known about the steps of invadopodia disassembly although the Rho GTPase RhoG and tyrosine phosphorylation of paxillin have been implicated in the process (Badowski et al., 2008; Goicoechea et al., 2017).

The dramatic rearrangement of the actin cytoskeleton during invadopodia formation is regulated by the Rho family of small GTPases (Murphy and Courtneidge, 2011). Rho GTPases function as molecular switches that cycle between an active GTP-bound and an inactive-GDP bound state. The activation of Rho proteins is catalyzed by RhoGEFs (guanine nucleotide exchange factors), whereas RhoGAPs (GTPase-activating proteins) mediate their inactivation. There are 80 RhoGEFs and 66 RhoGAPs in humans, allowing cells to tightly regulate the activity of Rho proteins through multiple pathways (Kreider-Letterman et al., 2022a; Rossman et al., 2005). Since RhoGAPs play a critical role in the termination of signal transduction, mutations in genes encoding RhoGAPs have drastic consequences and underlie several human diseases including cancer (Kreider-Letterman et al., 2022a). RhoGAPs are generally considered tumor suppressors, and the loss of GAP activity results in aberrant GTPase activity which can promote tumorigenesis (Kreider-Letterman et al., 2022a). However, compared with RhoGEFs, which have been studied more extensively, there is significantly less known about RhoGAPs, especially regarding their role in cancer progression and metastasis. The need to better characterize RhoGAPs in cancer progression includes invadopodia as there is increasing evidence that RhoGAPs play a role in their regulation (Al Haddad et al., 2020; Bravo-Cordero et al., 2011; Diring et al., 2019; Nakahara et al., 1998; Noll et al., 2019).

Several Rho proteins are known to be directly involved in invadopodia formation, including Cdc42, Rac1, RhoA, RhoC, and RhoG (Ayala et al., 2009; Bravo-Cordero et al., 2011; Goicoechea et al., 2017; Moshfegh et al., 2014; Sakurai-Yageta et al., 2008; Yamaguchi et al., 2005). However, the molecular mechanisms of their spatiotemporal activity at invadopodia, as well as the identity of their upstream regulators and downstream effectors remain poorly characterized. Here, we conducted a candidate shRNA screen to find RhoGAPs involved in invadopodia formation and identified ARHGAP17 (RICH1, Nadrin), a Cdc42-specific GAP and a negative regulator of invadopodia in triple negative breast cancer cells.

Using TIRF and STORM microscopy and live cell analysis of Cdc42 activity via biosensors, we found that during invadopodia assembly, ARHGAP17 localizes specifically to the ring region of invadopodia where it functions to restrict the activity of Cdc42 to the invadopodia core. Interestingly, invadopodia start to disassemble when ARHGAP17 translocates from the ring to the core where it inactivates Cdc42 and promotes invadopodia disassembly. We also show that this shift in localization is mediated by the interaction between ARHGAP17 and CIP4, a Cdc42 effector which localizes to the core of invadopodia and is important for invadopodia assembly. Altogether, our results identify ARHGAP17 at the center of the dynamic regulation of invadopodia assembly and disassembly and provide new insights into the molecular mechanisms controlling these processes.

## Results

### ARHGAP17 is a negative regulator of invadopodia in breast cancer cells

To characterize the role of RhoGAPs during invadopodia formation, we used the triple negative breast cancer (TNBC) cell line SUM159 as a model. SUM159 cells produce invadopodia that are highly dynamic and typically form in clusters under the nucleus or as bands at the leading edge (Fig. 1 A and Video 1) (Goicoechea et al., 2017). When grown in culture, the cells form invadopodia clusters spontaneously at low levels (≈1% cells) (Fig. S1 B). However, treatment with the phorbol ester Phorbol 12,13-dibutyrate (PDBu) stimulates the formation of large numbers of invadopodia, with about 30% of cells forming invadopodia after 30 minutes (Video 1) (Goicoechea et al., 2009; Goicoechea et al., 2017). To identify RhoGAPs that may regulate invadopodia formation, we performed an shRNA screen on a panel of candidate RhoGAPs in PDBu treated SUM159 cells (Fig. 1 B). We tested 19 RhoGAP candidates using 5 targeting sequences per gene. Several RhoGAPs showed differences in the percentage of cells with invadopodia, typically an increase. Here, we focused on ARHGAP17, as silencing its expression showed the most prominent increases in the percentage of cells forming invadopodia, especially with shRNA sequences #3 and #5. We confirmed the efficiency of the shRNA targeting sequences by western blot analysis, which shows a significant reduction of ARHGAP17 expression in both sequences #3 and #5 when compared to the non-targeting control (Fig. 1 C). We also repeated the invadopodia assay with both shRNA sequences and confirmed that depletion of ARHGAP17 resulted in significant increases in the number of cells that form invadopodia after PDBu treatment (from 28.7% in CTRL cells to 79.3% and 72.3% in shRNAs #3 and #5 respectively) (Fig. 1 D). We also observed similar results using two unique CRISPR KO sequences for ARHGAP17 (Fig. S1 A and B). In addition, we were able to rescue the effects of silencing ARHGAP17 on invadopodia by re-expressing an shRNA resistant myc-ARHGAP17 (Fig. 1 E-G). The increase in cells forming invadopodia observed in ARHGAP17 KD was independent of PDBu treatment as silencing ARHGAP17 also induced a significant increase in the number of spontaneous invadopodia in untreated cells (from 1.3% in CTRL cells to 7.8% in ARHGAP17 KO) (Fig. S2 A and B). These results validated the specificity of the phenotype and demonstrated that the effect of silencing ARHGAP17 expression is not dependent on PDBu treatment. Taken together our results suggest that ARHGAP17 functions as a negative regulator of invadopodia.

**Figure 1.**
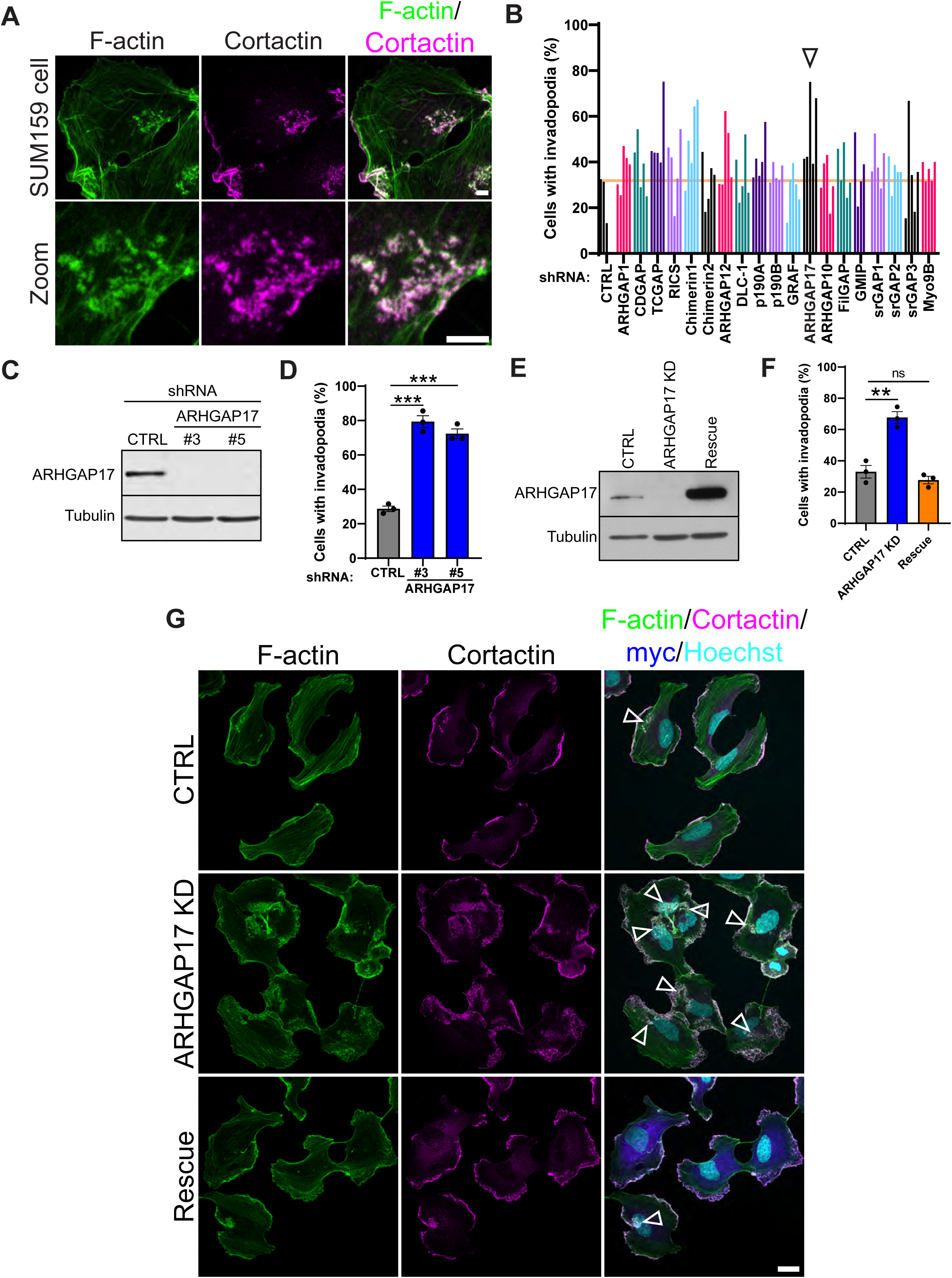
ARHGAP17 negatively regulates invadopodia. (**A**) Representative micrographs of a SUM159 cell with an invadopodia cluster. Cells were treated with PDBu for 30 minutes and stained for cortactin and F-actin (scale bar for both panels, 5 μm). (**B**) shRNA screen of candidate RhoGAPs in SUM159 cells. Cells stably expressing the indicated shRNAs were treated with PDBu for 30 minutes and stained for cortactin and F-actin as invadopodia markers. Colors represent different RhoGAPs, and individual bars represent unique shRNA targeting sequences. Results are expressed as the percentage of cells with invadopodia from 4 images per condition for a total of 7100 cells. (**C**) Western blot of ARHGAP17 expression in SUM159 cells stably expressing non-targeting and ARHGAP17-specific shRNAs. Tubulin was used as a loading control. (**D**) Percentage of cells with invadopodia in CTRL or ARHGAP17 KD SUM159 cells treated with PDBu for 30 minutes. Results are expressed as mean ± S.E.M. of 3 independent experiments in which at least 200 cells per condition per experiment were counted. ***, p<0.001. (**E**) Western blot of ARHGAP17 KD and Rescue with shRNA-resistant myc-ARHGAP17 in SUM159 cells. (**F**) Percentage of cells with invadopodia in CTRL, ARHGAP17 KD, and Rescue SUM159 cells treated PDBu for 30 minutes. Results are expressed as mean ± S.E.M. of 3 independent experiments in which at least 200 cells per condition per experiment were counted. **, p<0.01; ns, not significant. (**G**) Representative micrographs of CTRL, ARHGAP17 KD, and Rescue with myc-ARHGAP17 in SUM159 cells. F-actin and cortactin staining were used as markers for invadopodia while myc staining was used to visualize myc-ARHGAP17. Cells were treated with PDBu for 30 minutes before fixation and staining. Arrowheads highlight invadopodia clusters (scale bar, 20 μm).

### ARHGAP17 regulates invadopodia-based matrix degradation and inhibits invasion

To determine whether the increases in invadopodia in the absence of ARHGAP17 resulted in more invadopodia activity, we used a matrix degradation assay in which we plated SUM159 cells on a fluorescent gelatin/collagen IV matrix and treated them with PDBu for 24 hours (Martin et al., 2012). SUM159 cells degraded the matrix with a punctate pattern typical for invadopodia; characterized by F-actin puncta overlapping with a loss of matrix fluorescence, provided the cell had not moved before fixation (Fig. 2 A) (Artym et al., 2006). We quantified invadopodia activity in CTRL, ARHGAP17 KO, and Rescue cells by measuring the area of degradation/cell area. We found a significant increase in matrix degradation in ARHGAP17 KO cells (∼3-fold) which was rescued to CTRL levels by re-expressing ARHGAP17 (Fig. 2 B and C). These results demonstrate that the additional invadopodia formed in the absence of ARHGAP17 are functional.

**Figure 2.**
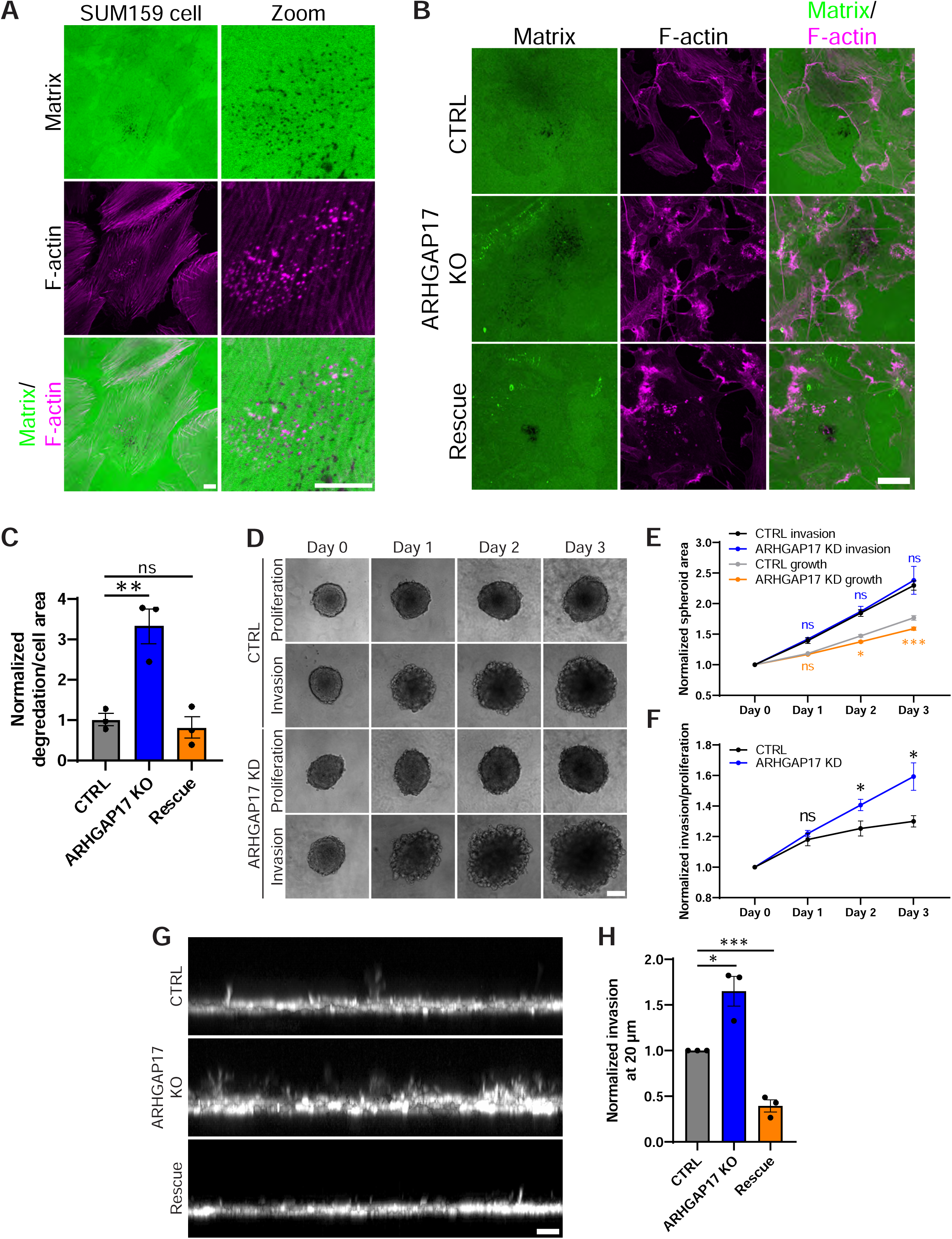
Loss of ARHGAP17 expression promotes invasion. (**A**) Representative micrographs of SUM159 cells degrading a fluorescent-gelatin and collagen IV substrate. F-actin staining was used as a marker for invadopodia. The zoomed panel on the right shows a cluster of invadopodia colocalizing with degradation spots (scale bar for left and right panels, 10 μm). (**B**) Representative micrographs of F-actin-stained SUM159 cells degrading fluorescent-gelatin and collagen IV matrix after 24-hour PDBu treatment (scale bar, 20 μm). (**C**) Quantification of matrix degradation by SUM159 cells. Results are expressed as mean ± S.E.M. of 3 independent experiments in which at least 20 random fields-of-view (FOV) per condition per experiment were measured. **, p<0.01; ns, not significant. (**D**) Micrographs of SUM159 spheroids grown with Matrigel to promote invasion and without Matrigel to observe spheroid growth (scale bar, 200 μm). (**E and F**) Quantification of spheroid size and quantification of invasion corrected for proliferation. Results are expressed as mean ± S.E.M. of 4 independent experiments in which at least 5 spheroids per condition per experiment were measured. *, p<0.05; ***, p<0.001; ns, not significant. (**G**) Z-view of inverse invasion assay of SUM159 cells expressing Scarlet-CAAX with brightest point projection in the y-axis (scale bar, 20 μm). (**H**) Quantification of area invaded by cells in the inverse invasion assay. Invasion was measured at 20 μm above the coverslip. Results are expressed as mean ± S.E.M. of 3 independent experiments in which at least 3 FOV per condition per experiment were measured. *, p<0.05; ***, p<0.001.

Since silencing the expression of ARHGAP17 results in increased matrix degradation via invadopodia, we would predict the cells to be more invasive when grown in 3D. To determine if ARHGAP17 KD cells are more invasive, we performed a 3D spheroid invasion assay (Vinci et al., 2015). In this assay, SUM159 cells were grown in ultra-low attachment plates that resulted in the formation of a spheroid reminiscent of a small tumor (Fig. 2 D). When the spheroids are embedded in Matrigel, invasive cells migrate away from the spheroid and into the matrix. Spheroid growth and invasion can then be quantified by measuring the area of the spheroid over time. We found that even though ARHGAP17 KD spheroids grew at a slower rate (Fig. 2 E), they were significantly more invasive when invasion was normalized by average proliferation (Fig. 2 F). To validate the spheroid invasion results without the effect of growth impacting the results we also performed an inverse invasion assay. In this assay, we plated SUM159 cells stably expressing mScarlet-CAAX (membrane marker) at the bottom of a plate, overlaid them with a layer of Matrigel and stimulated them to invade upwards by adding serum-containing media on top of the Matrigel (Fig. 2 G and Video 2). Quantification showed that ARHGAP17 KO cells invaded significantly more than the CTRL and that this effect was rescued by re-expressing ARHGAP17 (Fig. 2 H).

### ARHGAP17 regulates invadopodia dynamics

The dramatic increase in invadopodia observed in ARHGAP17 KD cells suggested that ARHGAP17 could be modulating invadopodia turnover. Interestingly, in the absence of ARHGAP17, not only were more cells able to form invadopodia compared to CTRL cells, but also the invadopodia clusters were larger (Fig. 3 A). This may reflect a change in invadopodia dynamics, possibly due to an increase in invadopodia lifetime or defects in disassembly signaling. To further explore this possibility, we performed a time-course experiment in which we treated SUM159 cells (CTRL, ARHGAP17 KD, and Rescue) with PDBu and quantified the percentage of cells forming invadopodia at successive time points after treatment (Fig. 3 B). As we have previously described (Goicoechea et al., 2017), CTRL cells showed a rapid spike in invadopodia forming cells at the 10-minute mark that then decreased rapidly until reaching equilibrium at 30 minutes, with approximately 30% of cells being positive for invadopodia at any given time afterwards. Interestingly, the initial increase in invadopodia was larger in ARHGAP17 KD cells (>80% positive cells at 10 min), and the decrease not as pronounced, with invadopodia percentages reaching an equilibrium at a much higher percentage (>60% after 30 min). Re-expressing ARHGAP17 restored the normal time-course dynamics (Fig. 3 B). When we calculated the difference in percentage of cells with invadopodia between the 10- and 30-min time points, i.e., the drop-off from peak to equilibrium, the difference was significantly smaller in ARHGAP17 KD cells when compared to CTRL or Rescue (50% reduction in CTRL vs. 20% reduction in ARHGAP17 KD) (Fig. S3 A). These results suggest that in the absence of ARHGAP17, the invadopodia may be more stable and thus longer lived.

**Figure 3.**
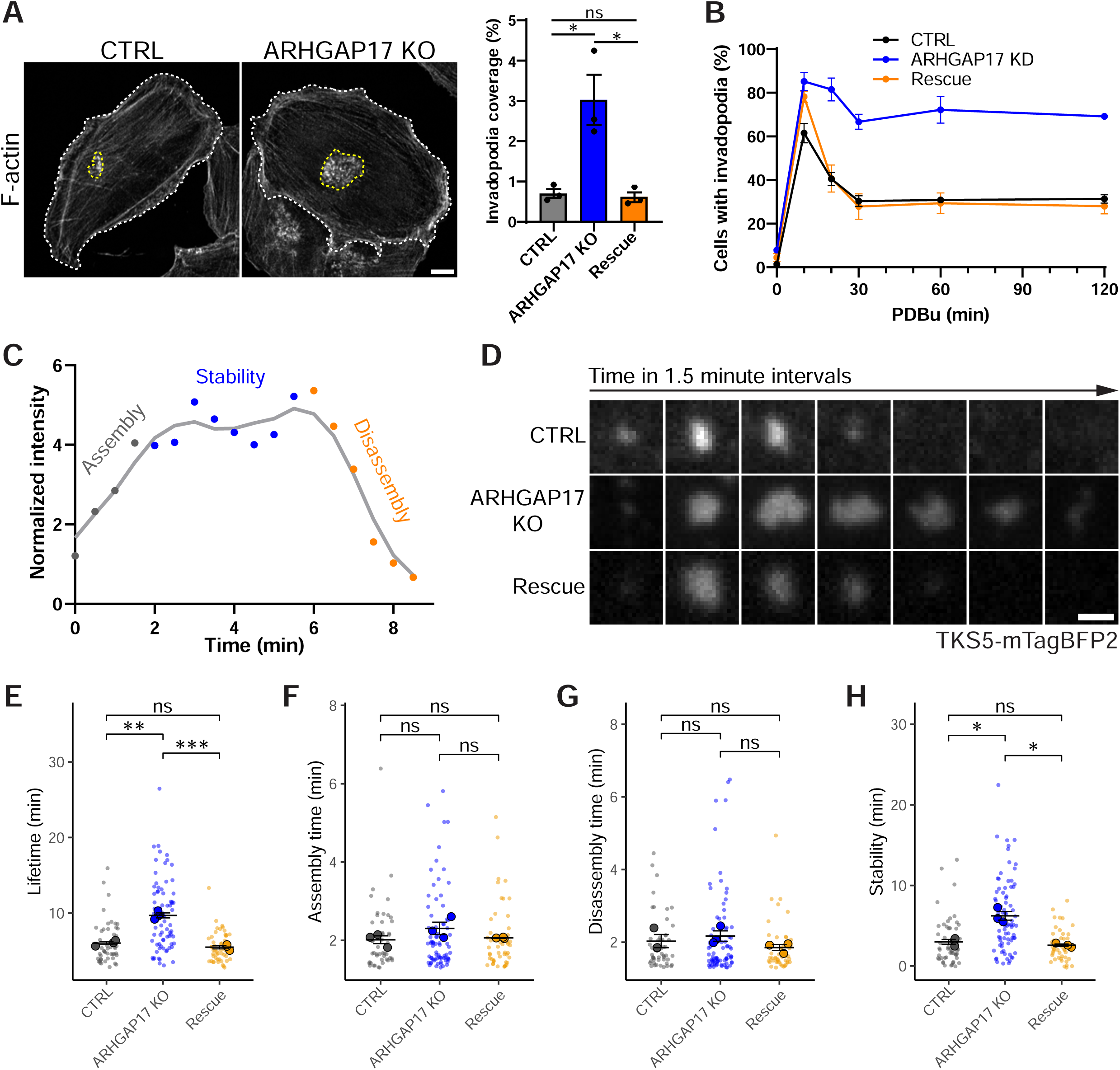
ARHGAP17 knockdown increases invadopodia stability and accumulation. (**A**) Representative micrographs of SUM159 cells stained for F-actin (scale bar, 10 μm) and quantification of percent invadopodia coverage after 30 min PDBu treatment. White dashed lines represent the cell outline and yellow dashed lines represent invadopodia clusters. Results are expressed as mean ± S.E.M. of 3 independent experiments in which at least 30 cells per condition per experiment were counted. *, p<0.05; ns, not significant. (**B**) Time course of percentage of SUM159 cells forming invadopodia after PDBu treatment at different intervals. Results are expressed as mean ± S.E.M. of 3 independent experiments in which at least 200 cells per condition per experiment were counted. (**C**) Example of single invadopodia dynamics showing assembly, stability, and disassembly phases as indicated by measuring TKS5-mTagBFP2 intensity over time at 30 second intervals after PDBu treatment. The gray line represents a second-order smoothing polynomial. (**D**) Representative micrographs of single invadopodia dynamics of SUM159 cells expressing TKS5-mTagBFP2 as a marker for invadopodia after PDBu treatment (scale bar, 1 μm). (**E-H**) Quantification of invadopodia dynamics. Results are expressed as mean ± S.E.M. of 3 independent experiments in which at least 12 invadopodia per condition per experiment were measured. Large circles represent biological replicates, small circles represent technical replicates. *, p<0.05; **, p<0.01; ***, p<0.001; ns, not significant.

We next wanted to characterize the role of ARHGAP17 on invadopodia dynamics. The dynamics of single invadopodia can be divided into three defined stages: assembly, stability, and disassembly. This is illustrated in Fig. 3 C, in which the intensity of the invadopodia marker TKS5-mTagBFP2 was measured over the lifetime of a single invadopodia. Using live Total Internal Reflection Fluorescence (TIRF) microscopy and a custom algorithm inspired by a similar method previously used to quantify focal adhesions (Berginski et al., 2011), we were able to accurately measure the dynamics of invadopodia formation after PDBu treatment with high enough resolution to resolve the individual invadopodia from the clusters (Fig. 3 D). Using this method, we found that individual invadopodia have increased overall lifetime in ARHGAP17 KO cells (Fig. 3 D and E), and that this increase in lifetime was a result of an increase in the duration of the stability phase (Fig. 3 H) and not due to significant changes in the duration of the assembly or disassembly phases (Fig. 3 F and G). We also confirmed that the effect of silencing ARHGAP17 on lifetime was independent of PDBu treatment, as we observed a similar increase in lifetime in spontaneous invadopodia in the absence of ARHGAP17 (Fig. S4 A). Interestingly, we found no significant changes in the assembly or disassembly rates (Fig. S4 B and C). Finally, we analyzed the results using a Principal Component Analysis (PCA) which confirmed that ARHGAP17 KO invadopodia form a distinct population based on lifetime and stability with only minimal impact from the other variables (Fig. S4 D). These results demonstrate that ARHGAP17 regulates the dynamics of invadopodia and that, in the absence of ARHGAP17, invadopodia are more stable which is associated with increased accumulation of invadopodia within clusters.

### ARHGAP17 is targeted to invadopodia through multiple domains

To visualize the localization of endogenous ARHGAP17 in cells we generated a polyclonal antibody against a small peptide from the C-terminus of ARHGAP17. The antibody recognized a single band by western blot, which disappeared upon KD or KO of ARHGAP17 (Fig. 1 C and Fig. S1 A). In PDBu-treated SUM159 cells, the ARHGAP17 antibody showed a clear signal at the invadopodia clusters. This signal was lost in ARHGAP17 KO cells confirming the specificity of the antibody (Fig. 4 A). A similar result was observed when expressing an ARHGAP17-GFP construct in ARHGAP17 KO SUM159 cells, with ARHGAP17-GFP targeting efficiently to invadopodia as marked by TKS5-mTagBFP2 (Fig. 4 B). Live imaging analysis showed that ARHGAP17-GFP is recruited early during invadopodia formation, almost simultaneously with TKS5-mTagBFP2, and remains associated until the invadopodia disappears (Fig. 4 B and Video 3). Interestingly, we observed that ARHGAP17 did not perfectly colocalize with TKS5 and had a more diffuse appearance with the signal localized in some cases adjacent to, or in the periphery of TKS5 puncta (Fig. 4 B, zoom panel).

**Figure 4.**
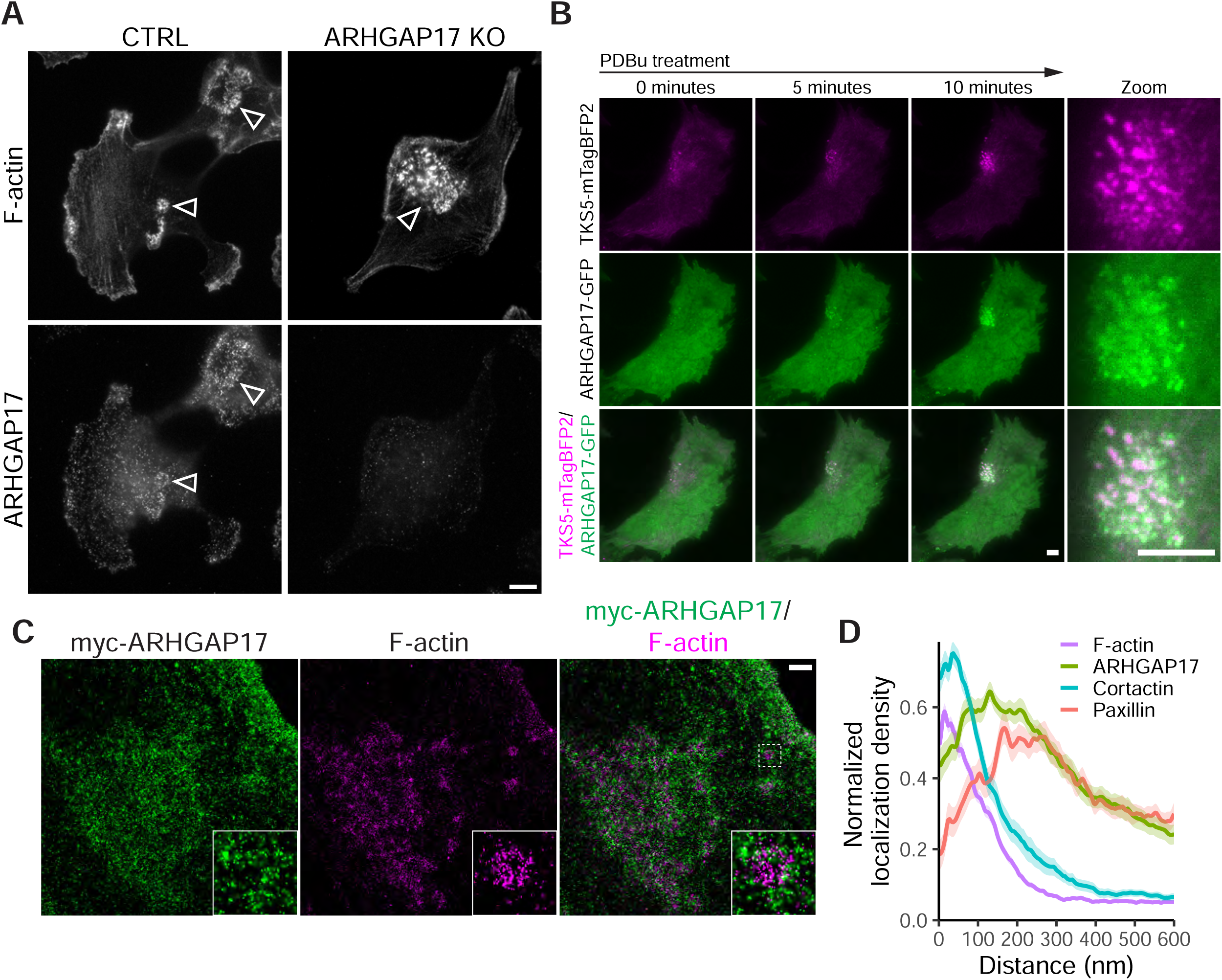
ARHGAP17 localizes to invadopodia. (**A**) Representative micrographs of CTRL and ARHGAP17 KO SUM159 cells stained for F-actin and ARHGAP17. Arrowheads highlight invadopodia clusters (scale bar, 10 μm). (**B**) Representative TIRF micrographs of a SUM159 cell expressing TKS5-mTagBFP2 as a marker for invadopodia and ARHGAP17-GFP (expressed in ARHGAP17 KO cells) forming an invadopodia cluster after 30 min PDBu treatment (scale bar for both panels, 5 μm). (**C**) STORM reconstructions of SUM159 cells treated with PDBu showing myc-ARHGAP17 (expressed in ARHGAP17 KO cells) localization to invadopodia as defined by F-actin staining (scale bar, 1 μm). (**D**) Quantification of STORM reconstructions using the radial intensity analysis of F-actin, myc-ARHGAP17, cortactin, and paxillin in invadopodia. Lines are expressed as mean ± S.E.M. of 3 independent experiments in which at least 20 invadopodia per condition per experiment were measured.

Invadopodia are typically organized with a core region comprised of actin, actin nucleators, and scaffolding proteins, and a surrounding ring comprised of adhesion and adapter proteins such as paxillin and vinculin (Branch et al., 2012). Our initial results suggested that ARHGAP17 may be localized preferentially to the ring region of invadopodia. To better define the localization of ARHGAP17 at invadopodia, we utilized STORM (Stochastic Optical Reconstruction Microscopy) reconstructions of invadopodia using F-actin (phalloidin staining) and cortactin as markers for the invadopodia core, and paxillin as a marker for the ring. We then compared the intensity distribution of these proteins to that of myc-ARHGAP17. We used ARHGAP17 KO SUM159 cells stably expressing myc-ARHGAP17 to facilitate the use of anti-myc antibodies, which provided better results with STORM than staining for the endogenous protein. From these results, ARHGAP17 appears to share its localization pattern more with paxillin than actin or cortactin (Fig. 4 C and Fig. S5 A and B).

To compare the localization of different invadopodia proteins we developed a semi-automatic macro script in ImageJ that measures the intensity of invadopodia proteins in relation to the center of the invadopodia (hereafter referred to as radial intensity analysis). In this script, a line is drawn across the center of each invadopodia. The line is then sequentially rotated 120 times at 3° intervals and the intensity profile is measured at each of the intervals for all fluorescent channels. The mean of all measurements is then used to display the average intensity related to the distance from the center of the invadopodia (Fig. 4 D). When this method is applied to the STORM reconstructions, we found that ARHGAP17 localization extends further away from the center of the invadopodia when compared to the core proteins cortactin and F-actin, and aligns closely with the ring protein paxillin, suggesting it is a ring protein (Fig. 4 D). However, unlike paxillin, a fraction of ARHGAP17 is also localized to the core. Fig. 4 D shows that the intensity of myc-ARHGAP17 remains high at the core when compared to that of paxillin, which decreases sharply at the invadopodia center (Fig. 4 C and Fig. S5 A and B). An equivalent localization of ARHGAP17 to the ring was also observed in the spontaneous invadopodia that form in non-treated cells (Fig. S6 A and B).

To determine how ARHGAP17 is targeted to invadopodia, we generated and expressed a series of domain deletion mutants in ARHGAP17 KO SUM159 cells (Fig. 5A and Fig. S7 A). Using TIRF microscopy for its high signal-to-noise ratio, we found that both the N-BAR domain and the C-terminus Proline-rich region are important for targeting ARHGAP17 to invadopodia (Fig. 5 B-E). We then used the radial intensity analysis to precisely define the localization for each of the mutants (Fig. 5 C). Our results show that upon deletion of the C-terminus (ΔC), ARHGAP17 localization decreases at the core, but remains associated with the ring, which suggests that the Proline-rich region is important for targeting ARHGAP17 to the center of invadopodia (Fig. 5 B and C). In this mutant, the ring diameter is also wider (Fig. 5 D). In contrast, when the N-BAR domain is deleted (ΔBAR), there is an increase in the localization of ARHGAP17 to the core, which is accompanied by a slight reduction in the ring and a more diffuse localization pattern (Fig. 5 B-D). Interestingly, the N-BAR domain alone is sufficient to target to the core of invadopodia, where it is distributed in a punctate pattern (Fig. 5 B-E). Deletion of either the C-terminus or the N-BAR domain reduced the total amount of ARHGAP17 recruited to invadopodia compared to the WT protein, which suggests that targeting to invadopodia is partially compromised in both mutants (Fig. 5 E).

**Figure 5.**
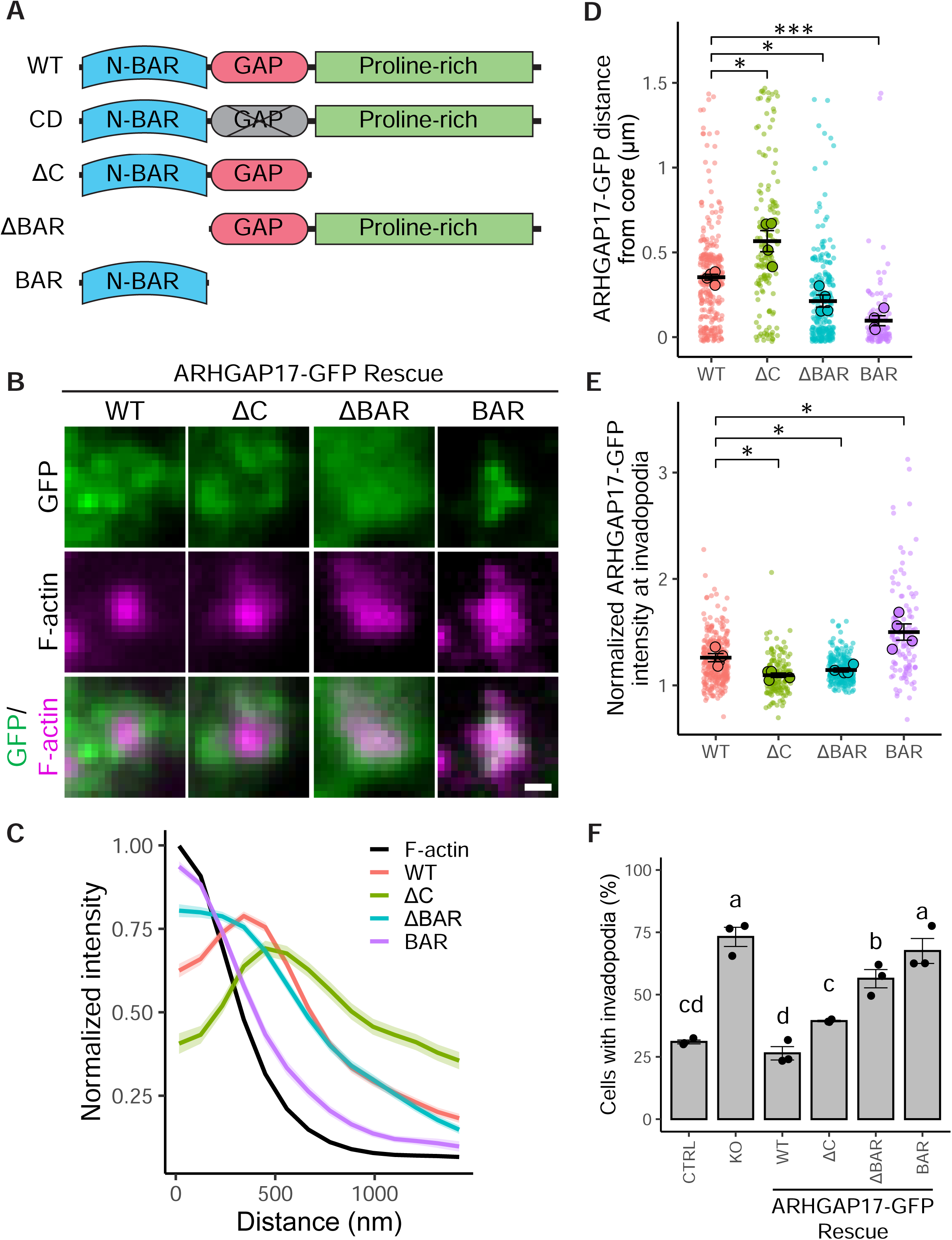
ARHGAP17 targets invadopodia through both the N-BAR and proline-rich domains. (**A**) Domain structure of ARHGAP17 and the ARHGAP17 mutants used in this study. (**B**) Representative TIRF micrographs of individual invadopodia in SUM159 cells expressing ARHGAP17-GFP domain deletion mutants in ARHGAP17 KO cells. Cells were treated with PDBu for 30 min and stained for F-actin as a marker for invadopodia (scale bar, 0.5 μm). (**C**) Radial intensity analysis of F-actin and ARHGAP17-GFP mutants in invadopodia. Lines are expressed as mean ± S.E.M. of 4 independent experiments in which at least 26 invadopodia per condition per experiment were measured. (**D and E**) Quantification of ARHGAP17 distance from the center of invadopodia based on peak intensity and quantification of normalized intensity of ARHGAP17 at invadopodia. Results are expressed as mean ± S.E.M. of 4 independent experiments in which at least 26 invadopodia per condition per experiment were measured. Large circles represent biological replicates, small circles represent technical replicates. *, p<0.05; ***, p<0.001. (**F**) Percentage of cells with invadopodia in CTRL, ARHGAP17 KO, and Rescue with ARHGAP17 domain deletion mutants after 30 minutes PDBu treatment. Results are expressed as mean ± S.E.M. of 3 independent experiments in which at least 200 cells per condition per experiment were counted. Statistical analysis is a 1-way ANOVA with a Tukey’s post hoc test. Different letters indicate significant differences between conditions as defined by p<0.05.

We also tested these deletion mutants for their ability to rescue the ARHGAP17 KO phenotype. Our results show that the ΔC mutant rescues the levels of invadopodia efficiently to a level that is close but still significantly different than that of the WT Rescue. The ΔBAR mutant also showed a reproducible level of rescue although it was less efficient than the ΔC mutant. In contrast, the N-BAR domain alone showed no evidence of rescue, which confirms that the rest of the protein is not only important for targeting but also for ARHGAP17’s function. (Fig. 5 F). From these results, we conclude that ARHGAP17 regulates invadopodia by its ability to associate to both the ring and core structures through the N-BAR domain and C-terminus, respectively.

### ARHGAP17 regulation of invadopodia requires its RhoGAP activity for Cdc42

*In vitro* assays have initially shown ARHGAP17 to have RhoGAP specificity for all three of the ubiquitous Rho GTPases: Cdc42, Rac1, and RhoA (Harada et al., 2000). However, later studies suggest it is specific for Cdc42 and Rac1 *in vitro* with preference for Cdc42 in cells (Richnau and Aspenstrom, 2001; Richnau et al., 2004; Wells et al., 2006). To assess the specificity of endogenous ARHGAP17 in SUM159 cells, we performed a GAP pulldown assay using glutathione S-transferase (GST)-tagged, constitutively active mutants of RhoA, Rac1, and Cdc42 (Fig. 6 A) (García-Mata et al., 2006). We found that ARHGAP17 precipitated with active Cdc42 but not with RhoA or Rac1. Furthermore, when the expression of ARHGAP17 was silenced in SUM159 cells, there was a significant increase in total Cdc42 activity in the cells as determined by an affinity pulldown assay (Fig. 6 B). From these results, we determined that in SUM159 cells ARHGAP17 is specific for Cdc42.

**Figure 6.**
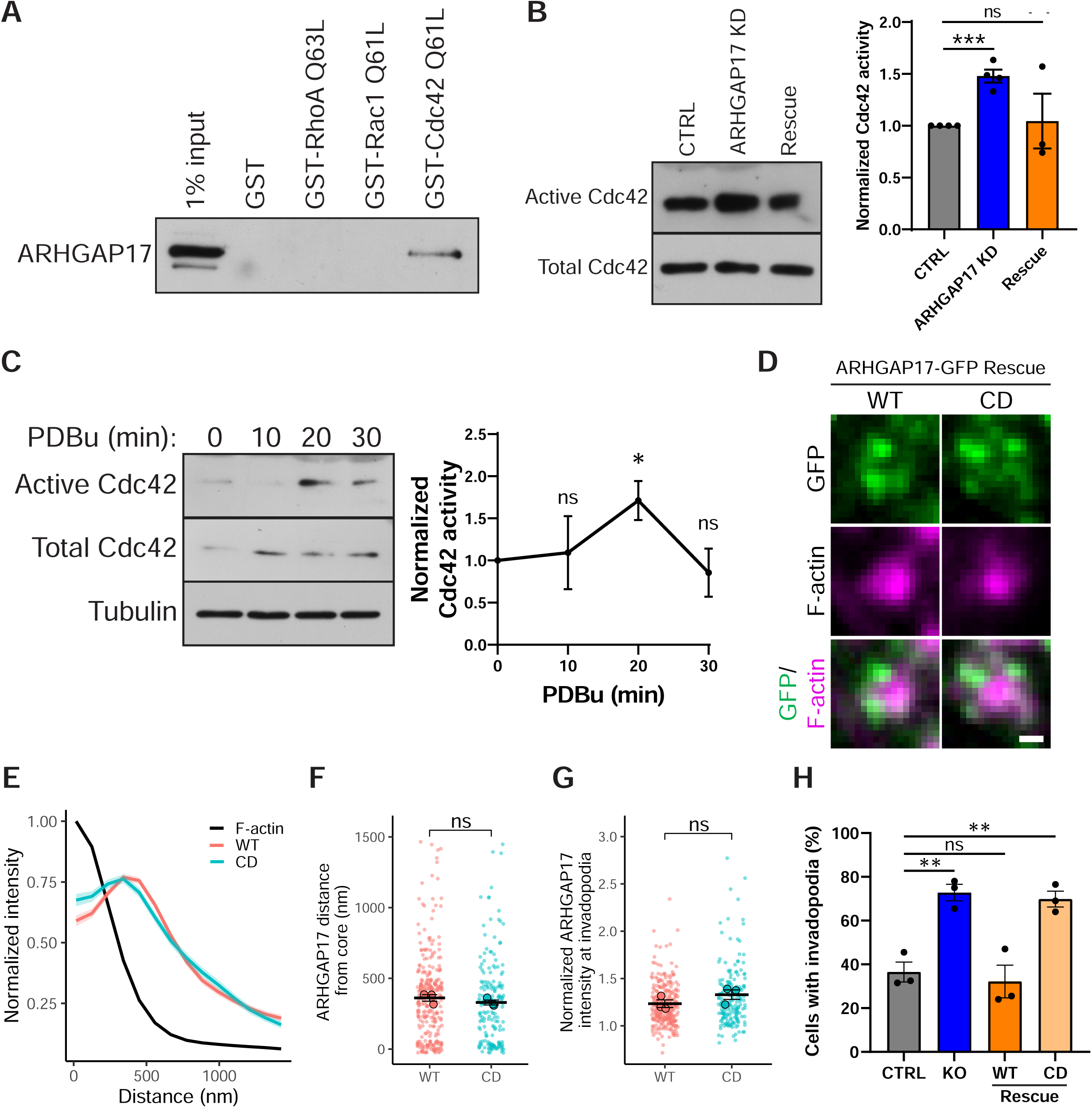
ARHGAP17 Cdc42-specific catalytic activity is required for its function at invadopodia. (**A**) Pulldown of endogenous ARHGAP17 in SUM159 cells using constitutively active RhoA (Q63L), Rac1 (Q61L), and Cdc42 (Q61L) tagged with Glutathione S-transferase (GST) or with GST alone. (**B**) Pulldown of active Cdc42 using the p21 binding domain (PBD) of p21 activated kinase 1 protein (PAK). Results are expressed as mean ± S.E.M. of 4 independent experiments. ***, p<0.001; ns, not significant. (**C**) PBD pulldown of active Cdc42 in SUM159 cells after treatment with PDBu at the indicated time points. Left panel shows a representative WB; right panel shows the quantification of 3 independent experiments. Results are expressed as mean ± S.E.M. *, p<0.05; ns, not significant. (**D**) Representative TIRF micrographs of ARHGAP17 KO SUM159 cells forming single invadopodia after PDBu treatment stained for F-actin as a marker for invadopodia and expressing either WT or catalytic dead (CD) ARHGAP17-GFP (expressed in ARHGAP17 KO background) (scale bar, 0.5 μm). (**E**) Radial intensity analysis of F-actin, WT ARHGAP17-GFP, and CD ARHGAP17-GFP in invadopodia. Lines are expressed as mean ± S.E.M. of 3 independent experiments in which at least 40 invadopodia per condition per experiment were measured. (**F and G**) Quantification of ARHGAP17-GFP distance from the center of invadopodia based on peak intensity and normalized intensity of ARHGAP17 at invadopodia. Results are expressed as mean ± S.E.M. of 3 independent experiments in which at least 40 invadopodia per condition per experiment were measured. Large circles represent biological replicates, small circles represent technical replicates. ns, not significant. (**H**) Percentage of cells with invadopodia in CTRL, ARHGAP17 KO, and Rescue with either WT or CD ARHGAP17-GFP after 30 minutes PDBu treatment. Results are expressed as mean ± S.E.M. of 3 independent experiments in which at least 200 cells per condition per experiment were counted. **, p<0.01; ns, not significant.

Since Cdc42 signaling is known to be essential for the formation of invadopodia (Ayala et al., 2009; Di Martino et al., 2014; Goicoechea et al., 2014; Sakurai-Yageta et al., 2008; Yamaguchi et al., 2005; Zagryazhskaya-Masson et al., 2020), we performed a time course assay in which we measured Cdc42 activity in SUM159 cells at 10-minute intervals following PDBu treatment (Fig. 6 C). Our results showed a peak in Cdc42 activity in CTRL cells that roughly correlates with the time course of invadopodia formation (see Fig. 3 B). ARHGAP17 KD cells also showed an increase in Cdc42 activity at the same time point (20 min). However, the Cdc42 increase was not as prominent as in CTRL cells, likely because the basal activity of Cdc42 is already comparatively higher at time zero in the ARGHAP17 KD cells (Fig. 6 B and Fig. S8 A, compare the activity between t0 and t20). These results suggest that ARHGAP17 modulates the activity of Cdc42 during the formation of invadopodia.

To test this prediction, we mutated the critical Arginine finger motif in the RhoGAP domain of ARHGAP17 (R288A), which renders it catalytically dead (CD) (Harada et al., 2000). We then quantified both the localization of ARHGAP17 CD to invadopodia and its ability to rescue the ARHGAP17 KO phenotype. Our results demonstrated that the catalytic activity is not required for targeting ARHGAP17 to invadopodia, as the CD mutant localization and radial intensity analysis were indistinguishable from that of the WT protein (Fig. 6 D and E and Fig. S9 A). The distance from the core and the intensity at the invadopodia were also not significantly different between WT and CD ARHGAP17 (Fig. 6 F and G). However, despite targeting correctly, the CD mutant was unable to rescue the KO phenotype, which suggests the RhoGAP activity is required for ARHGAP17 to regulate invadopodia turnover (Fig. 6 H).

### Spatiotemporal regulation of Cdc42 by ARHGAP17 at invadopodia

Since ARHGAP17 regulation of invadopodia requires its Cdc42-specific RhoGAP activity, we wanted to characterize the spatiotemporal regulation of Cdc42 activity and ARHGAP17 at invadopodia. We initially analyzed the dynamics of Cdc42 in live cells using a previously described Cdc42-FRET biosensor, which we co-expressed in SUM159 cells with mCherry-cortactin as an invadopodia marker (Reinhard et al., 2017). This unimolecular FRET sensor, and similar designs for Rac1 and RhoA, have been previously characterized (Kedziora et al., 2016; Reinhard et al., 2017; Timmerman et al., 2015; van Unen et al., 2016). The Cdc42 FRET biosensor behaved as previously described, showing higher Cdc42-activity at the edge of dynamic cells protrusions, which are enriched in mCherry-cortactin (Fig. S10 A) (Machacek et al., 2009). During invadopodia formation, again visualized with mCherry-cortactin, we observed an increase in Cdc42 activity at invadopodia (Fig. S10 A). The FRET signal increased at invadopodia as they assembled and decreased during their disassembly until returning to background levels (Fig. S10 B). To determine whether we could map the FRET signal representing the activity to Cdc42 to the core or ring of each invadopodia puncta, we applied the radial intensity analysis to compare the FRET biosensor signal to the signal of mCherry-cortactin (core marker). Our results showed that the FRET signal broadly overlaps with that of cortactin, suggesting the activity is mostly restricted to the core (Fig. S10 D). However, the spatial resolution of the FRET sensor is significantly lower than that of mCherry-cortactin, and the variability between individual puncta is high, so it was difficult to definitively conclude that Cdc42 was only active at the core. To obtain higher resolution images of the dynamics of Cdc42 activity in live cells, we took advantage of a localization based Cdc42 biosensor (dTomato-wGBD) that has been recently optimized for imaging in mammalian cells. The sensor consists of the GTPase-binding domain of the Cdc42 effector WASP (wGBD) linked to the dTomato fluorescent protein (Fig. S11 A). The sensor encodes the Cdc42 binding domain of WASP only, so it does not interact with other WASP-binding proteins such as Cdc42 effector proteins (e.g., CIP4), so the accumulation of signal in cells should reflect its association with active Cdc42 pools (Tian et al., 2000). This sensor design has been extensively used in the past, particularly in Xenopus studies, but was typically not sensitive enough for mammalian cell studies (Benink and Bement, 2005; Kim et al., 2000; Vaughan et al., 2011). Our new Cdc42 sensor has been designed using the same iterative approach we recently described for a RhoA sensor (Mahlandt et al., 2021). To verify the specificity of the sensor in SUM159 cells, we used an assay that measures the translocation of the sensor to the nucleus when co-expressed with different nuclear localized constitutively active Rho GTPases as we have previously described (Mahlandt et al., 2021). Our results show that the dTomato-wGBD sensor only relocates to the nucleus in the presence of constitutively active Cdc42, suggesting it is highly specific for Cdc42 (Fig. S11 B). Quantification of nuclear over cytosolic shows a >4-fold ratio increases when nuclear Cdc42 is expressed vs. the empty nuclear control. Nuclear Rac1 showed only residual binding, while RhoA showed no detectable binding (Fig. S11 B and C). Similarly to what has been reported when Cdc42 activity was measured using FRET sensors, the wGBD sensor, but not the empty control sensor, localizes to the leading edge of protruding lamellipodia, and its signal decreases upon retraction (Fig. S11 D and E and Video 4) (Machacek et al., 2009).

We then tested the localization of the sensor in SUM159 cells stimulated with PDBu to form invadopodia. Upon treatment of PDBu, the sensor strongly re-localizes to newly forming invadopodia, indicating an increase in Cdc42 activity specifically at invadopodia (Fig. 7 A and Video 5). In contrast, the negative control showed no accumulation at invadopodia clusters (Fig. S12 A and Video 6). We also followed the activity of Cdc42 in single invadopodia over time and compared it with the intensity of ARGHAP17-GFP and TKS5-mTagBFP2. We found that the sensor signal follows the progression of the invadopodia, increasing in intensity during its assembly and decreasing as the invadopodia disappears (Fig. 7 B and Video 7). Interestingly, we observed that Cdc42 activity appears early during the initial stages of invadopodia formation, overlapping closely with the pattern of TKS5. In contrast, ARHGAP17-GFP appears later during the assembly phase (∼1 min later than TKS5/wGBD in the example shown in Fig. 7B), and it also shows a slight increase in signal intensity during invadopodia disassembly (Fig. 7 B and Video 7). Quantitative analysis of intensity over time in multiple invadopodia confirmed this observation (Fig. 7 C). Cross-correlation analysis between TKS5-mTagBFP2, ARHGAP17-GFP, and dTomato-wGBD confirmed these observations, and showed that there is a positive correlation between the three proteins, with the strongest correlation observed between Cdc42 activity and TKS5 (Fig. 7 D). The analysis shows that the activity of Cdc42 occurred slightly earlier in time than TKS5. In contrast, the correlation between ARHGAP17 and Cdc42 activity or TKS5 was weaker, most likely due to the ring localization of ARHGAP17, with ARHGAP17 appearing later than both Cdc42 and TKS5. In addition, our results also showed that in the absence of ARHGAP17 (ARHGAP17 KO) there was a significant increase in the maximum Cdc42 activity at invadopodia when compared to CTRL or ARHGAP17 Rescue cells, confirming that ARHGAP17 plays a role in the local regulation of Cdc42 at invadopodia (Fig. 7 F).

**Figure 7.**
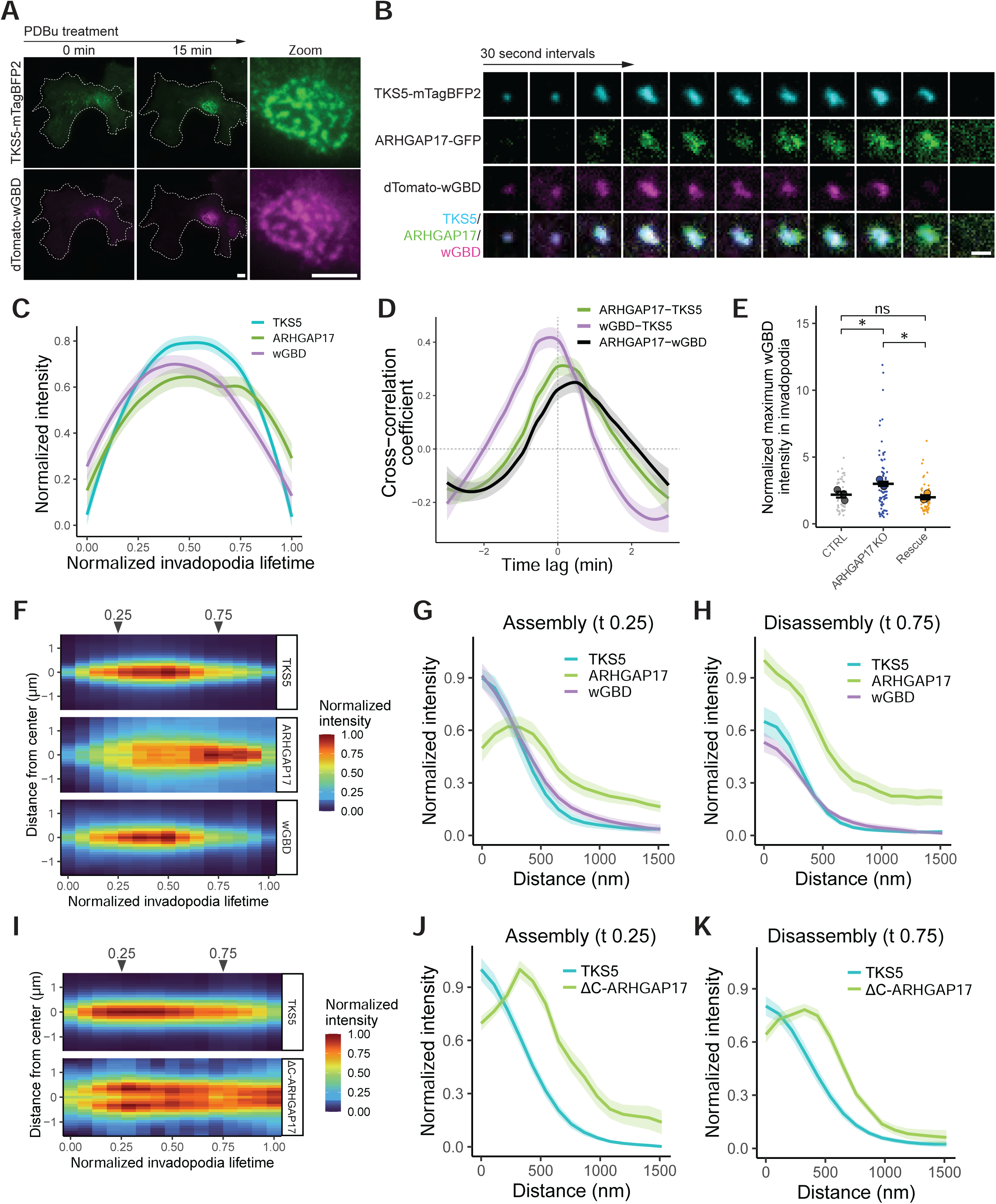
ARHGAP17 regulates the spatiotemporal dynamics of Cdc42 activity during invadopodia turnover. (**A**) Representative TIRF micrographs of a SUM159 cell expressing TKS5-mTagBFP2 as a marker for invadopodia and dTomato-wGBD as a marker for Cdc42 activity and forming an invadopodia cluster after PDBu treatment (scale bar for both panels, 5 μm). (**B**) Representative TIRF micrographs of a SUM159 cell forming an individual invadopodia after PDBu treatment. Cells were expressing TKS5-mTagBFP2 as a marker for invadopodia, ARHGAP17-GFP (expressed in ARHGAP17 KO cells), and dTomato-wGBD as a marker for Cdc42 activity (scale bar, 1 μm). (**C**) Quantification of TKS5-mTagBFP2, ARHGAP17-GFP (expressed in ARHGAP17 KO cells), and dTomato-wGBD intensity at individual invadopodia. Intensity and time have been rescaled between values of 0 and 1 for each invadopodia to allow for comparison. Lines are loess smoothing and 95% confidence from 3 independent experiments in which at least 10 invadopodia per condition per experiment were measured. (**D**) Cross-correlation analysis of TKS5-mTagBFP2, ARHGAP17-GFP (expressed in ARHGAP17 KO background), and dTomato-wGBD intensity. Higher values denote positive correlation and lag denotes the time shift effect on correlation between signals. Lines are loess smoothing and 95% confidence from 3 independent experiments in which at least 10 invadopodia per condition per experiment were measured. (**E**) Quantification of the peak dTomato-wGBD intensity at individual invadopodia after PDBu treatment, normalized to the average whole-cell intensity before treatment. Results are expressed as mean ± S.E.M. of 3 independent experiments. Large circles represent biological replicates, small circles represent technical replicates. *, p<0.05; ns, not significant. (**F**) Radial intensity profiles of individual invadopodia from live cells expressing TKS5-mTagBFP2, ARHGAP17-GFP (expressed in ARHGAP17 KO cells), and dTomato-wGBD plotted over time. Time scales between invadopodia are rescaled between 0 and 1 to allow for alignment of intensity values. Results are expressed as the mean of 8 representative invadopodia. (**G and H**) Radial profile plots taken from time 0.25 and 0.75 of panel (**F**) showing relative intensity and localization of the invadopodia markers while the invadopodia are assembling and disassembling, respectively. Results are expressed as mean ± S.E.M. of 8 invadopodia. (**I**) Radial intensity profiles of individual invadopodia from live cells expressing TKS5-mTagBFP2 and ΔC-ARHGAP17-GFP (expressed in ARHGAP17 KO cells) plotted over time. Time scales between invadopodia are rescaled between 0 and 1 to allow for alignment of intensity values. Results are expressed as the mean of 13 representative invadopodia. (**J and K**) Radial profile plots taken from time 0.25 and 0.75 of panel (**I**) showing relative intensity and localization of the invadopodia markers while the invadopodia are assembling and disassembling, respectively. Results are expressed as mean ± S.E.M. of 13 invadopodia.

To characterize the spatiotemporal dynamics of ARHGAP17 and Cdc42 throughout the lifecycle of invadopodia, we used the radial intensity analysis for each frame in time-lapse series of single invadopodia co-expressing TKS5-mTagBFP2, ARHGAP17-GFP, and the wGBD Cdc42 localization sensor (as in Fig. 7 B). A representative kymograph of several invadopodia averaged together is shown, displaying the signal intensity over time for all the markers analyzed (x-axis, normalized to be comparable), as well as the localization of the signal in relation to the center of the invadopodia (y-axis) (Fig. 7 F). These results showed that Cdc42 activity (wGBD) was tightly localized at invadopodia, specifically at the core throughout the lifetime of invadopodia. During the invadopodia assembly phase, ARHGAP17 is recruited preferentially to the invadopodia ring as expected from our results from fixed images in Fig. 4. Surprisingly, when the invadopodia starts to disassemble, the localization of ARHGAP17 shifts from the ring to the core, and this shift coincides with a decrease in Cdc42 signal at the core. This change in localization can also be visualized as line profiles of selected time points from the time series data (Fig. 7 G and H). At an early stage during invadopodia assembly (t 0.25), ARHGAP17 was found predominantly in the ring (Fig. 7 G), while later during invadopodia disassembly (t 0.75), ARHGAP17 was shifted to the core (Fig. 7 H). This shift of ARHGAP17 to the core is also associated with a decrease in the peak intensity of both TKS5 and wGBD signal (Fig. 7 H) when compared to the peak intensity during assembly (Fig. 7 G). When the kymograph analysis is repeated with the ΔC-ARHGAP17 mutant, which is impaired in core localization, the shift to the invadopodia core at disassembly is completely lost (Fig. 7 I-K). This shows that the shift in localization towards the core during disassembly is dependent on the C-terminus of ARHGAP17. Taken together, these data suggest that ARHGAP17 at the ring restricts the activity of Cdc42 to the core during invadopodia assembly and moves to the core later, where it inactivates Cdc42 to promote invadopodia disassembly.

### An ARHGAP17/CIP4/Cdc42 complex regulates invadopodia

We have shown that ARHGAP17 regulates Cdc42 at invadopodia in a RhoGAP-dependent manner. However, we do not know what Cdc42 effector is downstream of this signaling pathway. A potential candidate is Cdc42-interacting protein 4 (CIP4), a Cdc42 effector which has been previously shown to interact with ARHGAP17 (Richnau and Aspenstrom, 2001). Furthermore, CIP4 also targets to invadopodia and regulates their formation in breast cancer cells (Pichot et al., 2010). To validate these observations in our model, we generated SUM159 cells stably expressing shRNA for CIP4 (Fig. 8 A) and found that, as described previously (Pichot et al., 2010), there was a significant decrease in the percentage of cells forming invadopodia when the expression of CIP4 was silenced (Fig. 8 B and C). We then stained for endogenous CIP4 in PDBu-treated SUM159 cells and confirmed that CIP4 also targets specifically to invadopodia in these cells (Fig. 8 D) and localizes predominantly to the invadopodia core (Fig. 8 E). We also confirmed that ARHGAP17 binds CIP4 at endogenous levels using co-immunoprecipitation (Fig. 8 F) and that the proline-rich region of ARHGAP17 is required for CIP4 binding as was previously shown (Fig. 8 G) (Richnau and Aspenstrom, 2001). We next wanted to determine if ARHGAP17 activity is coupled with CIP4 as part of a complex to regulate Cdc42 activity as has been frequently observed with RhoGEFs and some RhoGAPs (Lawson and Ridley, 2018). Using GST-Cdc42 Q61L and CD ARHGAP17-GFP (which does not interact with Cdc42), we found that myc-CIP4 can bind to active Cdc42 and CD ARHGAP17-GFP simultaneously (Fig. 8 G). These results suggest that CIP/ARHGAP17/Cdc42 may form a complex in cells which could potentially coordinate the transition between activation and inactivation of Cdc42.

**Figure 8.**
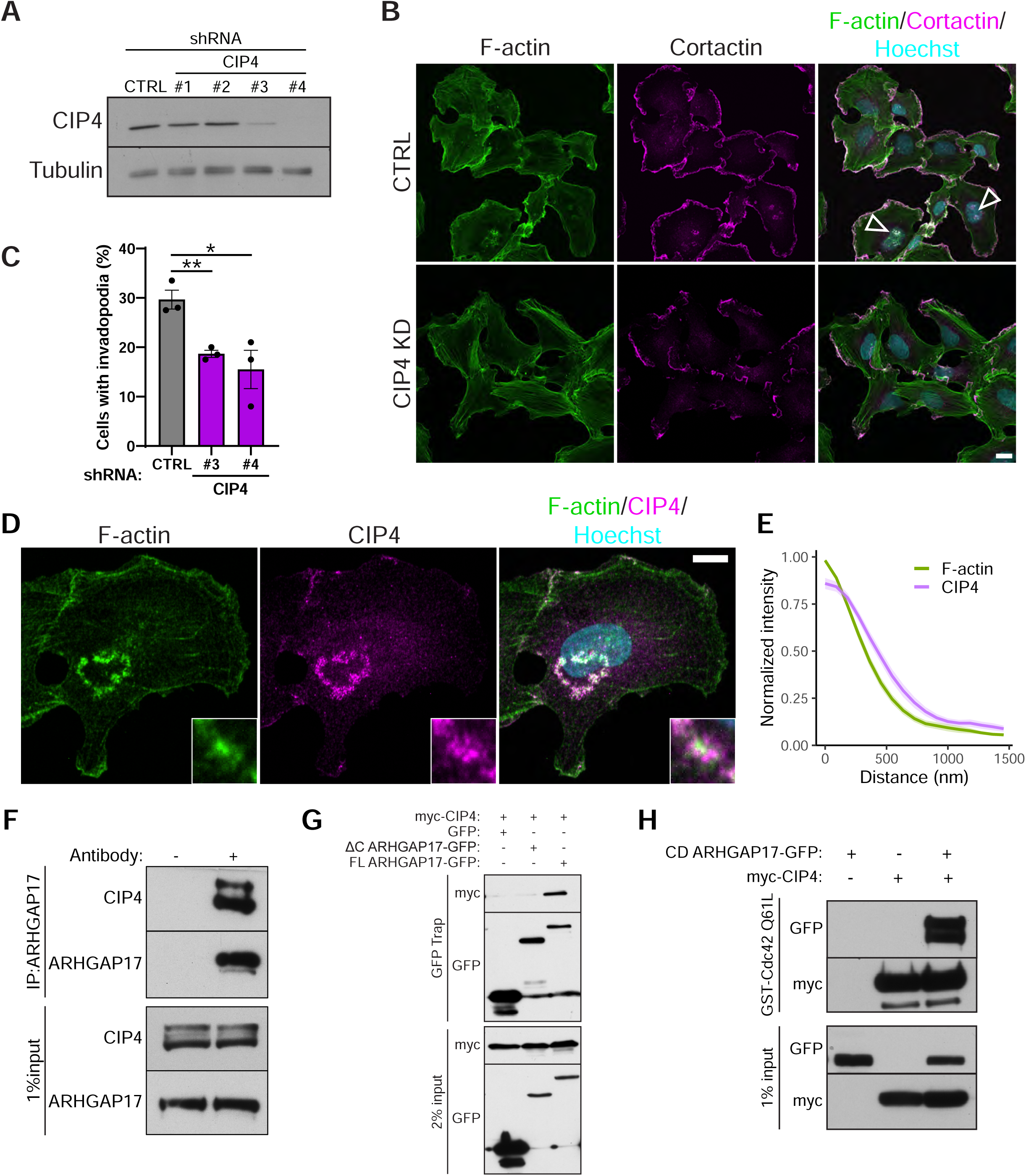
ARHGAP17 interacts with CIP4 to regulate invadopodia. (**A**) Representative western blot of CIP4 expression in SUM159 cells expressing non-targeting (CTRL) or CIP4-specific shRNAs. (**B**) Representative micrographs of CTRL and CIP4 KD (shRNA #4) SUM159 cells. F-actin and cortactin staining were used as markers for invadopodia. Cells were treated with PDBu for 30 minutes before fixation and staining. Arrowheads highlight invadopodia clusters (scale bar, 20 μm). (**C**) Percentage of cells with invadopodia in CTRL and CIP4 KD SUM159 cells treated PDBu for 30 minutes. Results are expressed as mean ± S.E.M. of 3 independent experiments in which at least 200 cells per condition per experiment were counted. *, p<0.05; **, p<0.01. (**D**) Representative micrographs of SUM159 cells treated with PDBu for 30 min and stained for F-actin and endogenous CIP4 (scale bar, 10 μm). (**E**) Radial intensity analysis of F-actin and CIP4 in invadopodia. Lines are expressed as mean ± S.E.M. of 3 independent experiments in which at least 10 invadopodia per condition per experiment were measured. (**F**) Western blot of co-immunoprecipitation of endogenous ARHGAP17 and CIP4 in SUM159 cells. (**G**) Western blot of co-immunoprecipitation of myc-CIP4 with GFP, ΔC ARHGAP17-GFP, or WT ARHGAP17-GFP. (**H**) Western blot of affinity precipitation of myc-CIP4 and CD ARHGAP17-GFP using GST-Cdc42-Q61L bound to Glutathione Sepharose 4B.

We then wanted to characterize the roles of the ARHGAP17, CIP4, and Cdc42 activity in invadopodia. When analyzing individual invadopodia expressing CIP4-GFP, TKS5-mTagBFP2, and dTomato-wGBD, we found that CIP4 is present at invadopodia throughout their lifetime and overlaps closely with TKS5, both spatially and temporally (Fig. 9 A and B and Video 8). Interestingly, Cdc42 activity, as determined by the dTomato-wGBD biosensor, was present earlier in invadopodia lifetime than CIP4, while CIP4 was still present in invadopodia after loss of Cdc42 activity (Fig. 9 A and B and Video 8). These temporal shifts between CIP4 and Cdc42 activity were confirmed using cross-correlation analysis (Fig. 9 C). To determine the role of the ARHGAP17-CIP4 interaction in the regulation of invadopodia, we investigated the localization of ARHGAP17 (ARHGAP17-GFP expressed in ARHGAP17 KO cells) in CTRL cells compared to cells overexpressing myc-CIP4. Interestingly, in cells overexpressing myc-CIP4, we observed a dramatic shift from the typical ARHGAP17 localization enriched at the ring, to a predominantly core localization (Fig. 9 D and E: top and middle panels). This was dependent on the interaction between ARHGAP17 and CIP4 since the ΔC ARHGAP17-GFP mutant, which does not interact with CIP4, failed to relocate to the core when co-expressed with myc-CIP4 (Fig. 9 D and E: bottom panels). Cross-correlation analysis of invadopodia clusters confirmed these results, with a high cross-correlation coefficient between ARHGAP17-GFP and myc-CIP4, which decreased drastically for the ΔC ARHGAP17-GFP and myc-CIP4 pair (Fig. 9 F), even though both proteins still localized to invadopodia (Fig. 9 D; bottom panels). These results suggest that the translocation of ARHGAP17 from the ring to the core observed upon initiation of invadopodia disassembly depends on ARHGAP17’s ability to bind to the core localized Cdc42 effector CIP4.

**Figure 9.**
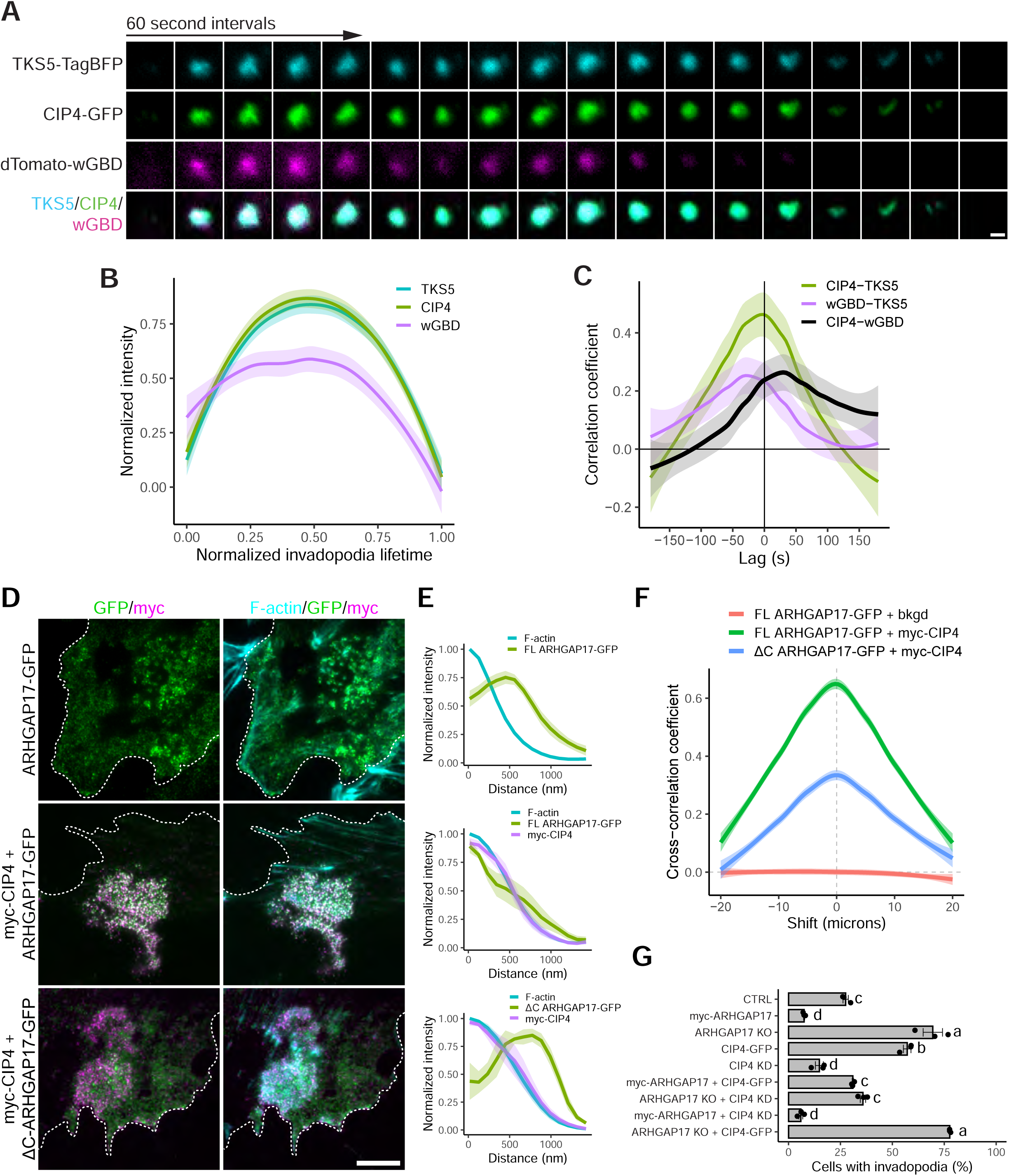
Spatiotemporal dynamics of CIP4 and ARHGAP17. (**A**) Representative TIRF micrographs of a SUM159 cell forming a single invadopodia after PDBu treatment expressing TKS5-mTagBFP2 as a marker for invadopodia, CIP4-GFP (expressed in CIP4 KD cells), and dTomato-wGBD as a marker for Cdc42 activity (scale bar, 0.5 μm). (**B**) Quantification of TKS5-mTagBFP2, CIP4-GFP (expressed in CIP4 KD cells), and dTomato-wGBD intensity at individual invadopodia. Intensity and time were rescaled between values of 0 and 1 for each invadopodia to allow for comparison. Lines are loess smoothing and 95% confidence from 3 independent experiments in which at least 10 invadopodia per condition per experiment were measured. (**C**) Cross-correlation analysis of TKS5-mTagBFP2, CIP4-GFP (expressed in CIP4 KD cells), and dTomato-wGBD intensity. Higher values denote positive correlation and lag denotes the time shift effect on correlation between signals. Lines are loess smoothing and 95% confidence from 3 independent experiments in which at least 10 invadopodia per condition per experiment were measured. (**D**) Representative TIRF micrographs of ARHGAP17 KO SUM159 cells transfected with the indicated constructs and treated with PDBu for 30 min. Cells were fixed and stained for F-actin and myc to detect myc-CIP4 (scale bar, 10 μm). (**E**) Radial intensity analysis of F-actin, ARHGAP17-GFP, and myc-CIP4 in invadopodia. Lines are expressed as mean ± S.E.M. of 3 independent experiments in which at least 10 invadopodia per condition per experiment were measured. (**F**) Van Steensil’s cross-correlation function of ARHGAP17-GFP colocalization with myc-CIP4 or the background signal of the anti-myc antibody at invadopodia clusters in SUM159 cells treated with PDBu for 30 min. Shift denotes the x-dimension movement of one channel in relation to the other. Lines are loess smoothing and 95% confidence from 3 independent experiments in which at least 10 invadopodia clusters per condition per experiment were measured. (**G**) Invadopodia assay of SUM159 cells after 30 minutes PDBu treatment in which CIP4 and ARHGAP17 were either silenced and/or overexpressed in all possible combinations as indicated in the graph. Results are expressed as mean ± S.E.M. of 3 independent experiments in which at least 200 cells per condition per experiment were counted. Statistical analysis is a 1-way ANOVA with a Tukey’s post hoc test. Different letters indicate significant differences between conditions as defined by p<0.05.

Finally, we wanted to determine how the expression levels of ARHGAP17 and CIP4 influence each other to regulate the invadopodia phenotype in SUM159 cells. To do this, we either silenced and/or overexpressed CIP4 and ARHGAP17 in all possible combinations and quantified the percentage of cells forming invadopodia after PDBu treatment. In agreement with what has been shown so far, our results showed that ARHGAP17 and CIP4 have antagonistic roles at invadopodia; CIP4 overexpression increased the number of cells with invadopodia while ARHGAP17 overexpression had an inhibitory effect. The opposite is true for silencing their expression, with ARHGAP17 KO increasing and CIP4 KD decreasing the number of cells with invadopodia (as shown in previous figures). When we tested combinations of overexpressing and silencing ARHGAP17/CIP4, the function of one protein either synergized or counteracted the effect of the other depending on the combination. If the two proteins were simultaneously overexpressed or silenced, the phenotypic effects balanced each other out. For example, the increase in invadopodia induced by CIP4 overexpression is restored to CTRL levels when ARHGAP17 is co-expressed. In contrast, a combination of silencing one gene and overexpressing the other worked to exacerbate the phenotype. For example, ARHGAP17 KO with CIP4 overexpression showed higher levels of invadopodia than either ARHGAP17 KO or CIP4-GFP overexpression alone (Fig. 9 G). Taken together, our results suggest that the expression levels of ARHGAP17 and CIP4 help regulate the equilibrium between invadopodia assembly and disassembly, and that CIP4 is key in recruiting ARHGAP17 to core to inactivate Cdc42 and initiate invadopodia disassembly.

## Discussion

Here, we have identified a novel role for ARHGAP17 in TNBC invadopodia dynamics by restricting the activity of Cdc42 in time and space during invadopodia turnover. Our results demonstrate that the function of ARHGAP17 is important throughout the invadopodia lifecycle. At early stages, during invadopodia growth, ARHGAP17 localizes to the invadopodia ring, which helps to limit the activity of Cdc42 to the actin core. Later, when invadopodia begin to disassemble, ARHGAP17 relocates to the core, where it inactivates Cdc42 and promotes the disassembly of invadopodia. We also show that this shift in localization is mediated by the interaction between ARHGAP17 and CIP4, a Cdc42 effector which localizes to the core of invadopodia and is important for invadopodia assembly (Fig. 10). It should be noted that, even in ARHGAP17 KO cells, invadopodia are still able to disassemble, and both the assembly and disassembly rate remain unchanged. The fact that they still assemble and disassemble at a similar rate may be indicative of redundancy in the system. ARHGAP17 has two closely related members of the RhoGAP family, ARHGAP44 (RICH2) and SH3BP1, which may be compensating for the loss of ARHGAP17 (Aspenstrom, 2018). One way to explain the increased stability then, is that the events that trigger disassembly, that is the recruitment of a Cdc42-GAP to the core, are either less frequent or delayed in the absence of ARHGAP17, and the redundant function of SH3BP1 and/or ARHGAP44 cannot completely compensate for this loss, and results in invadopodia remaining stable for a longer period. We plan to examine the role of the three family members in future studies.

**Figure 10.**
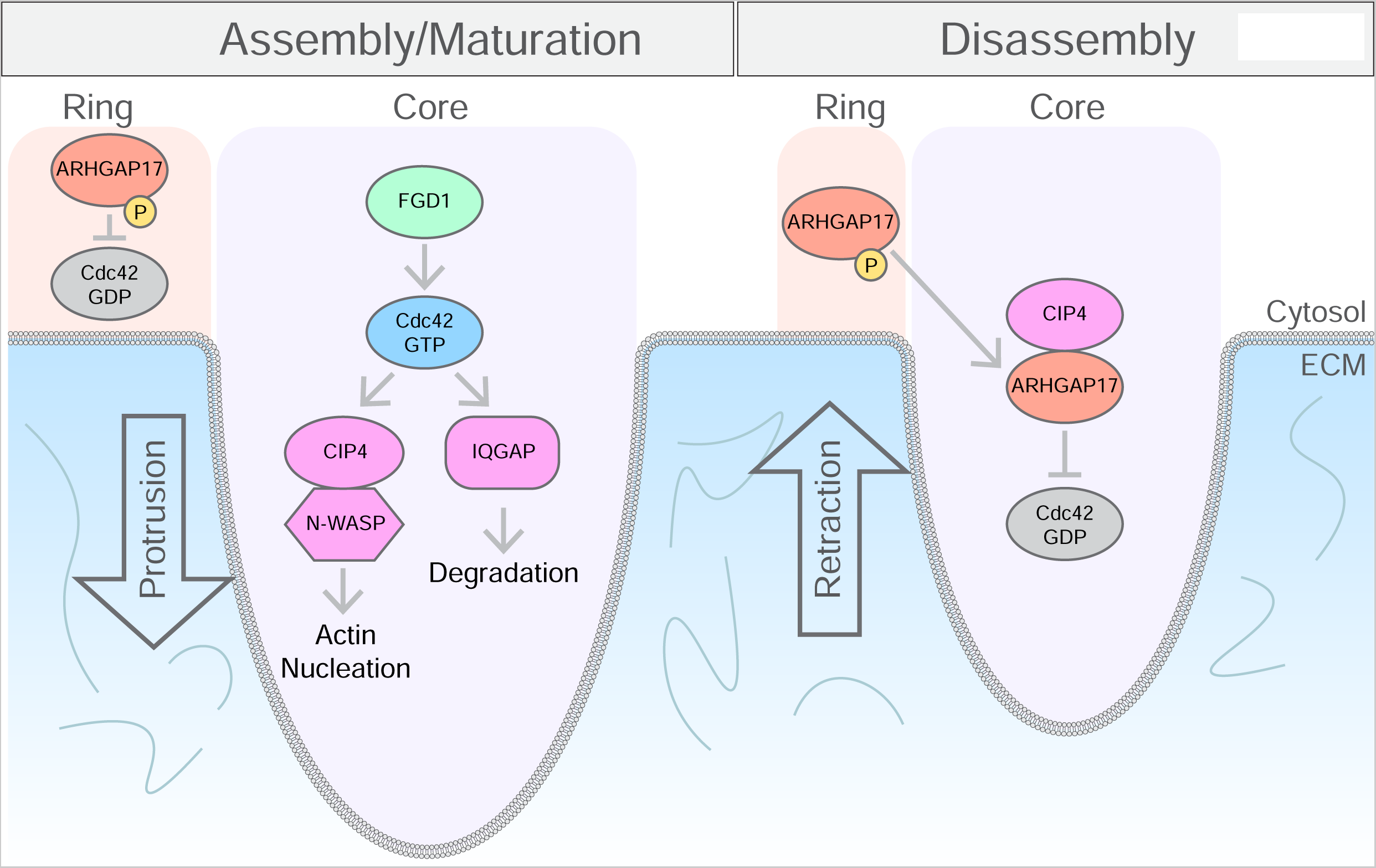
Model of ARHGAP17 function in invadopodia. During invadopodia assembly/maturation, ARHGAP17 is present at the invadopodia ring, which helps to restrict Cdc42 activity to the invadopodia core. Cdc42 is activated by a specific GEF at the invadopodia core, most likely FGD1, where it promotes actin nucleation through binding to N-WASP and CIP4 and stimulates degradation through its interaction with IQGAP. ARHGAP17 moves to the invadopodia core by binding to CIP4 where it inactivates Cdc42 to promote invadopodia disassembly.

Our report is the first to describe ARHGAP17 at invadopodia and a direct mechanism regarding the link between ARHGAP17 and cancer cell invasion. ARHGAP17 has been previously characterized as a tumor suppressor and found to be downregulated in a variety of cancers, including colon, breast, and cervical, where it functions to inhibit migration and invasion (Guo et al., 2019; Kiso et al., 2018; Pan et al., 2018; Pan et al., 2022; Tian et al., 2022). ARHGAP17 is downregulated by VEGF/NRP1 signaling in TNBC which was associated with increased Cdc42 activity, filopodia formation, and cell migration (Kiso et al., 2018). ARHGAP17 expression was also found to inhibit invasion and metastasis of colon cancer in association with decreased Wnt signaling (Pan et al., 2018), and in cervical cancer, ARHGAP17 suppressed cell proliferation and tumor growth by inhibiting PI3K/AKT signaling (Guo et al., 2019). Our results suggest that the tumor suppressor role of ARHGAP17 may depend in part in its ability to negatively regulate invadopodia by inactivating Cdc42, leading to a decreased invasive capacity.

Cdc42 activity has long been recognized to be essential for both invadopodia and podosome formation, regardless of the cell type or model system used (Ayala et al., 2009; Di Martino et al., 2014; Juin et al., 2014; Moreau et al., 2003; Yamaguchi et al., 2005). Using our improved localization-based sensor, we found that Cdc42 activity is present throughout the lifetime of invadopodia at the invadopodia core with the strongest activity being present at the assembly phase and a decrease in activity being associated with invadopodia disassembly. Cdc42 promotes the assembly of the actin core of invadopodia through its effector N-WASP, which induces actin nucleating by activation of the Arp2/3 complex (Desmarais et al., 2009; Juin et al., 2012; Juin et al., 2014; Lorenz et al., 2004; Monteiro et al., 2013; Yamaguchi et al., 2005; Yu et al., 2012). N-WASP also binds to the SH3 domain of the invadopodia protein, TKS5 (Oikawa et al., 2008). The link between Cdc42 and TKS5 is critical as Cdc42 activity and TKS5 are both essential for invadopodia formation and function (Di Martino et al., 2014). Cdc42 activity also promotes invadopodia function as marked by matrix degradation. Together with RhoA activity, Cdc42 has been shown to promote ECM degradation through its interaction with the Cdc42 effector, IQGAP (Sakurai-Yageta et al., 2008). Interactions between IQGAP and the exocyst protein complex control the delivery of matrix metalloproteases to the membrane of invadopodia (Sakurai-Yageta et al., 2008). Inactivation of Cdc42 at invadopodia by ARHGAP17 would then lead to the reduction of invadopodia formation and activity by shutting down Cdc42-mediated actin nucleation and metalloprotease transport.

The activities of several Rho GTPases have been associated with invadopodia formation and function, but very few have been studied at the level of detail shown in this manuscript. An exception is RhoC, which in contrast to what we observed for Cdc42, is active at the ring and inactive at the core. This is regulated by the combined action of the ring-localized p190 RhoGEF and the core localized p190RhoGAP (Bravo-Cordero et al., 2011). Whether p190RhoGAP shifts from the core-to-the ring to inactivate at RhoC is not clear from this study. However, like our study, it highlights the tight spatial and temporal regulation of Rho GTPases, even within invadopodia subdomains.

We have not characterized the RhoGEF responsible for the activation of Cdc42 in our system. A potential candidate is FGD1, which has been identified in previous reports as the RhoGEF that mediates the activation of Cdc42 at both podosomes and invadopodia (Ayala et al., 2009; Daubon et al., 2011; Zagryazhskaya-Masson et al., 2020). FGD1 localizes to invadopodia in metastatic melanoma and breast cancer cells through its interaction with TKS5 and promotes the assembly of the actin core in a Cdc42-dependent manner (Ayala et al., 2009; Zagryazhskaya-Masson et al., 2020). Another potential RhoGEF candidate is Vav1, which has also been found to play a role in invadopodia formation in pancreatic cancer cells by activating Cdc42 (Razidlo et al., 2014). In future studies, it will be interesting to characterize the RhoGEF in the ARHGAP17/Cdc42/CIP4 pathway and the interplay between GEF and GAP activity in regulating invadopodia dynamics.

Our study also shows that the Cdc42 effector CIP4 is important for the spatiotemporal regulation of ARHGAP17 at invadopodia and may play a key role in coordinating the timing of Cdc42 inactivation, as it can simultaneously interact with ARHGAP17 and Cdc42. We found that CIP4 localizes specifically to the invadopodia core, and that its interaction with ARHGAP17 is required for the ring-to-core translocation observed for ARHGAP17 at the initiation of invadopodia disassembly. CIP4 has previously been found to localize at invadopodia and function as a positive regulator of invadopodia formation through activation of N-WASP (Pichot et al., 2010). CIP4 seems to function as a scaffolding platform, as it can interact with Cdc42, N-WASP, Src, and ARHGAP17 (Dombrosky-Ferlan et al., 2003; Pichot et al., 2010; Richnau and Aspenstrom, 2001). CIP4 has two closely related family members which have also been shown to localize to invadopodia and promote their formation: FBP17 (Formin Binding Protein 17) and Toca-1 (transducer of Cdc42-dependent actin activity) (Chander et al., 2013; Suman et al., 2018; Yamamoto et al., 2011). FBP17 and Toca-1 both interact with Cdc42 and N-WASP, promote the activation of N-WASP, and stimulate actin nucleation through the Arp2/3 complex (Bu et al., 2010; Ho et al., 2004; Hu et al., 2011; Takano et al., 2008; Tsujita et al., 2013; Tsujita et al., 2006). Considering that silencing CIP4 expression does not completely abolish invadopodia formation, FBP17 and/or Toca-1 may compensate for CIP4 loss in SUM159 cells.

CIP4, FBP17, and Toca-1 can all activate N-WASP and have been shown to link its activity to membrane curvature sensing, membrane remodeling, and endocytosis through the F-BAR domains. (Chan Wah Hak et al., 2018; Hartig et al., 2009; Takano et al., 2008; Tsujita et al., 2006). It is possible that CIP4 regulates membrane curvature sensing and/or endocytosis at invadopodia, connecting these processes to actin polymerization through N-WASP activation. Interestingly, ARHGAP17 also encodes a BAR domain at the N-terminus, which is important for its targeting at invadopodia and may also play a role in curvature sensing and membrane remodeling/trafficking. What the functional connection between the BAR domains in CIP4 and ARHGAP17 remains to be determined.

It is not clear how the ring-to-core translocation is regulated, but phosphorylation is a possibility. Nagy and colleagues have shown that ARHGAP17 is phosphorylated by PKA/PKG at S702, and that this phosphorylation inhibits its binding to CIP4 (Nagy et al., 2015). It is possible then, that when targeted to the ring, ARHGAP17 is phosphorylated, and a yet to be characterized stimulus triggers its dephosphorylation at the initiation of disassembly, allowing it to bind to CIP4 and promoting the translocation to the core and subsequent inactivation of Cdc42. Interestingly, CIP4 is also phosphorylated by PKA at T225, and this phosphorylation appears to promote the formation of invadopodia and enhance its interaction with Cdc42 (Tonucci et al., 2019). Future work will shed light in the exact molecular mechanisms controlling the dynamics of ARHGAP17, Cdc42, and CIP4 in the formation of invadopodia, the role of the activators (RhoGEFs), and their effect on actin polymerization and other cellular functions.

## Materials and Methods

### Cell lines

HEK 293FT cells (Thermo Fisher Scientific) were used for all the pulldown experiments unless indicated otherwise. The SUM159 cells were a gift from Carol Otey (UNC-Chapel Hill, NC). The HEK 293FT cells were cultured in Dulbecco’s modified Eagle’s medium (DMEM) (GIBCO) containing 10% fetal bovine serum (FBS) (Sigma-Aldrich), and antibiotics (penicillin-streptomycin) (Corning). The SUM159 cells were cultured in Ham’s F12 (Cytiva) with 10% FBS, 5 μg/ml insulin (GIBCO), 1 μg/ml hydrocortisone (Sigma-Aldrich), and antibiotics. All cell lines were grown at 37 °C and 5% CO. Mycoplasma was tested regularly by staining with Hoechst 33342 (AnaSpec Inc.). Reagents and antibodies

### Reagents and antibodies

Antibodies against the following proteins were used: ARHGAP17 (Custom antibody, antigen sequence: EPHRSIFPEMHSDSASKDVPGR, Pacific Immunology); cortactin (sc-11408) (Santa Cruz); myc (13-2500) (Invitrogen); tubulin (T9028) and vinculin (V9131) (Sigma-Aldrich); CIP4 (612556), paxillin (610051), Cdc42 (610929), and Rac1 (610650) (BD Biosciences). F-actin was stained with Alexa Fluor-405, -488, - 594, -647, and -750 conjugated to phalloidin (Thermo Fisher Scientific). For immunofluorescence, the following secondary antibodies were used: Alexa Fluor-488, -568, -594, -647-conjugated anti-mouse-IgG and anti-rabbit-IgG secondary antibodies (A32723, A32731, A11004, A11011, A11032, A11037, A21236, A21245; Thermo Fisher Scientific). For western blot analysis the following secondary antibodies were used: horseradish peroxidase (HRP)conjugated anti-mouse-IgG and anti-rabbit-IgG (715-035-151 and 711-035-152; Jackson Immunoresearch). Phorbol-12,13-dibutyrate (PDBu) (P1269) was from Sigma-Aldrich.

### DNA constructs

Constructs generated for this study: pCMV-myc-ARHGAP17, pLenti-myc-ARHGAP17, pCI-CD-ARHGAP17-GFP, pC1-CMVdel-dTomato-1x-wGBD, pEGFP-ΔBAR-ARHGAP17, pLenti-BAR-ARHGAP17-GFP, pLenti-ΔC-ARHGAP17-GFP, pDG462-ARHGAP17-KO constructs were generated using the pDG462 backbone (Adikusuma et al., 2017) (exon 1 targets: CAGCTGGCTAACCAGACCG and GTTGAACTGCTTCTTCATGG; exon 3 targets: AGCATGGCACCGATGCCGAG and CCAAGCGCTTATGGGAATGG).

Previously published or purchased constructs: pAd-CMV-mCherry-Cortactin (Goicoechea et al., 2017); H2A-mTq2, H2A-mTq2-RhoA-G14V-ΔCAAX, H2A-mTq2-Rac1-G12V-ΔCAAX, and H2A-mTq2-Cdc42-G12V-ΔCAAX, pC1-CMVdel-dTomato (Mahlandt et al., 2021); pCMV6-myc-Trip10 (OriGene-MR208703); All shRNA were from the RNAi Consortium shRNA Library, purchased through Sigma-Aldrich (Table 1).

**Table 1.**
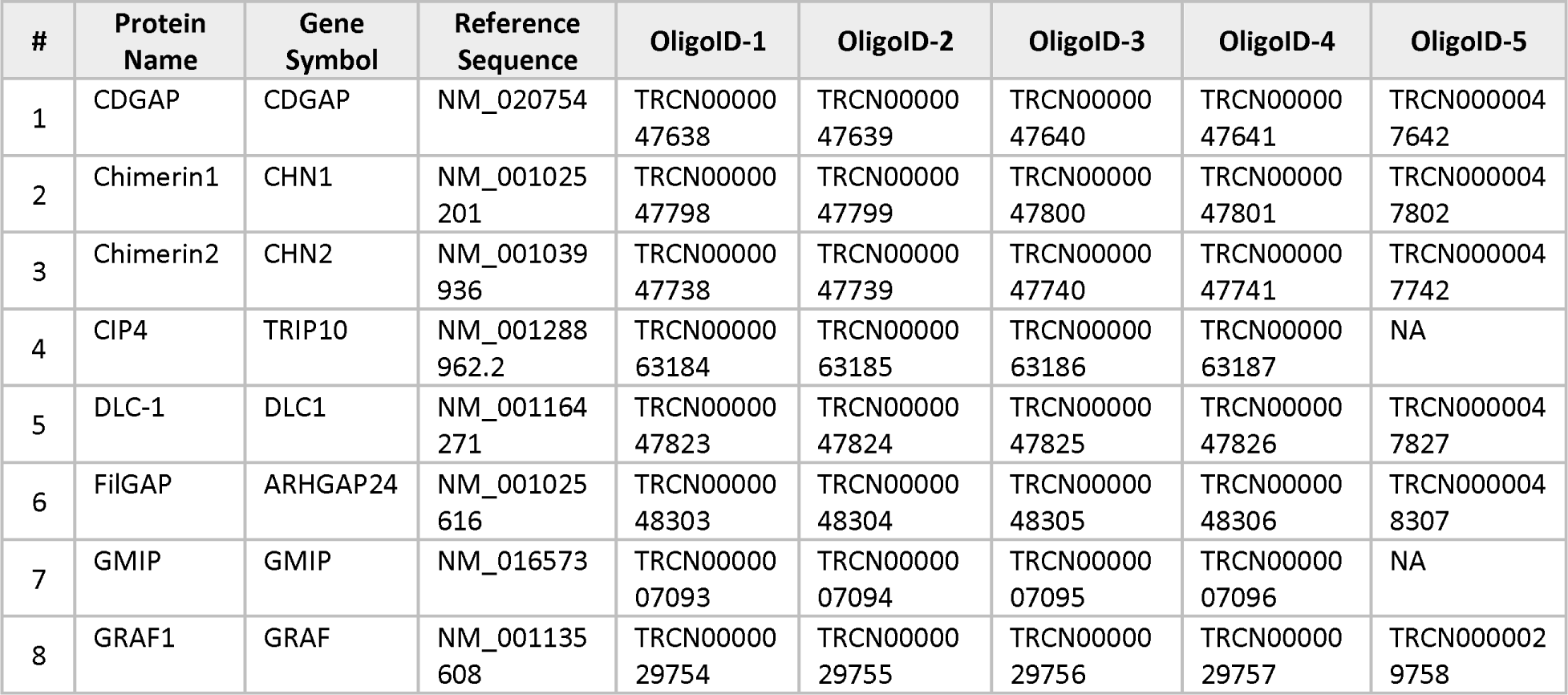
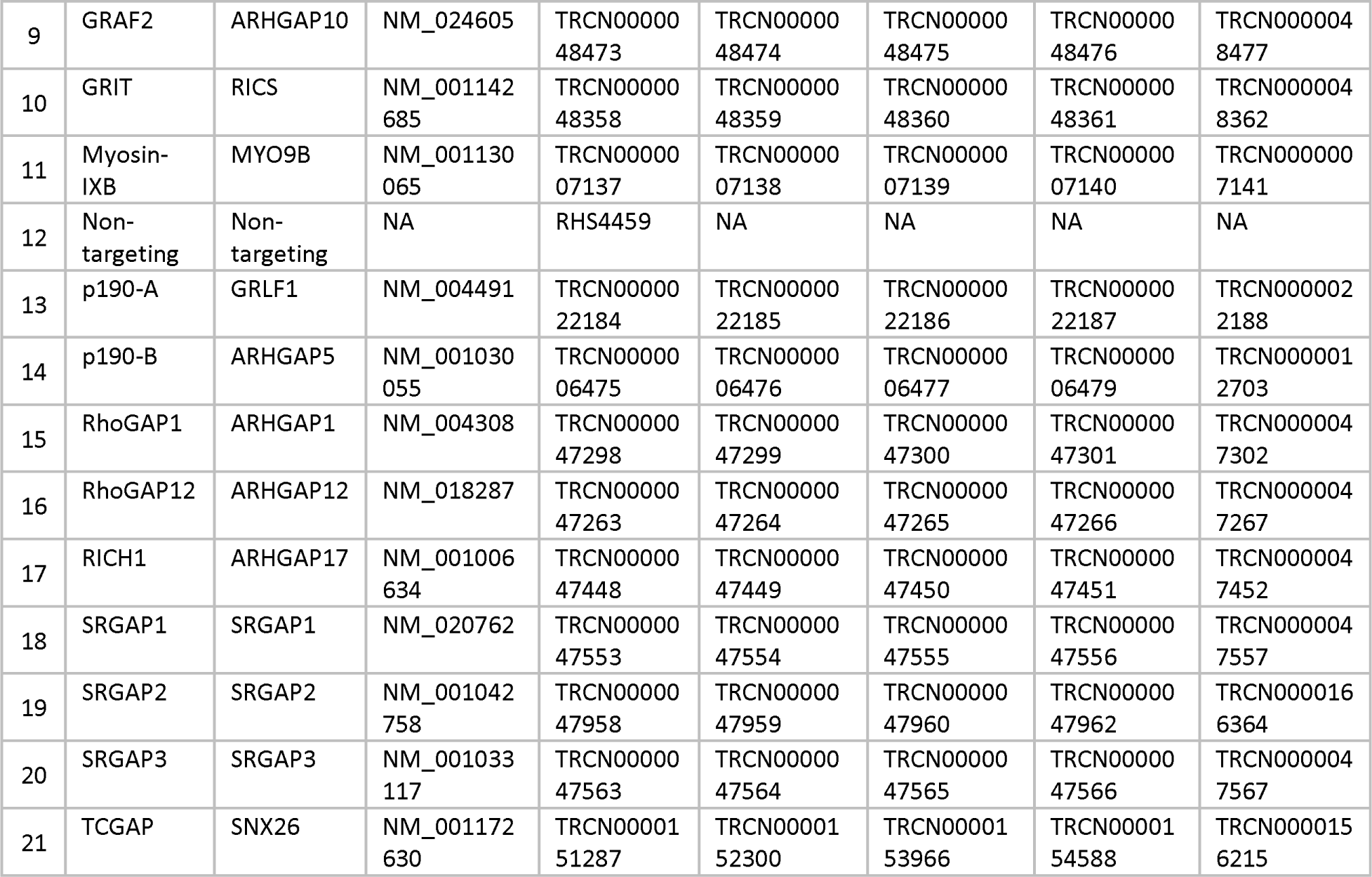
shRNAs used in this study

Constructs received as a gift: pCI-C-EGFP-Rich1 and pCI-C-EGFP-CIP4 (Harvey T. McMahon, MRC Laboratory of Molecular biology); pLV-mNeonGreen-CAAX and pLV-Scarlet-CAAX (Jaap van Buul, Sanquin); pmTagBFP2-N1-TKS5 and pmScarlet-N1-TKS5 (Lou Hodgson, Albert Einstein College of Medicine); pDG462 (Paul Thomas, Addgene plasmid # 100903); pFUGW-UbC-1T17 (Yi Wu, UConn Health, Farmington, CT) (Reinhard et al., 2017).

### Cell lysis and immunoblotting

Cells cultured on 100 mm tissue culture dishes were rinsed with TBS and 1 mM MgCl2 and then scraped into a lysis buffer containing 50 mM Tris-HCl pH 7.4, 10 mM MgCl2, 150 mM NaCl, 1% Triton X-100, and EZBlock protease inhibitor cocktail (BioVision). The supernatant was collected after centrifugation at 16,800g for 10 min. For immunoblotting, lysates were boiled in 2× Laemmli buffer, and were resolved by SDS-PAGE. The proteins were transferred onto PVDF and immunoblotted with the indicated antibodies. Immunocomplexes were visualized using the Immobilon Western Millipore Chemiluminescence HRP substrate (Millipore).

### Rho GTPase activity assay

Active Cdc42 pulldown experiments were performed as described previously (Goicoechea et al., 2014). Briefly, SUM159 cells were lysed in 50 mM Tris-HCl pH 7.4, 10 mM MgCl2, 150 mM NaCl, 1% Triton X-100, and EZBlock protease inhibitor cocktail. After clearing the lysates by centrifugation at 14,000 g for 5 min, the protein concentrations of the supernatants were determined, and equal amounts of total protein were incubated with 50 μg of GST–PBD, and rotated for 30 min at 4 °C. Subsequently the beads were washed three times in lysis buffer. Pull-downs and lysates were then immunoblotted for Cdc42.

### Immunofluorescence

For TIRF experiments, cells were grown in 8-well 1.5 glass bottom μ-slides (Ibidi), otherwise, cells were grown on 1.5 glass coverslips (Thermo Fisher Scientific). For immunofluorescence, cells were fixed for 10 min with 4% paraformaldehyde and quenched with 10 mM ammonium chloride. Cells were then permeabilized with 0.1% Triton X-100 in PBS for 7 min. The coverslips were then washed with PBS and blocked with PBS/2.5% goat serum/0.2% Tween 20 for 5 min, followed by 5 min of blocking with PBS/0.4% fish skin gelatin/0.2% Tween 20. Cells then were incubated with the primary antibody for 1 h at room temperature. Coverslips were washed 5 times with PBS/0.2% Tween 20 followed by 5 min of blocking with PBS/0.4% fish skin gelatin/0.2% Tween 20 and 5 min with PBS/2.5% goat serum/0.2% Tween 20. Secondary antibody diluted in the blocking solution was then added for 45 min, washed 5 times with PBS/0.2% Tween, and mounted on glass slides in Mowiol mounting solution (Electron Microscopy Sciences). For TIRF experiments, the cells were washed twice with PBS to remove any detergent and kept in PBS for imaging.

### STORM microscopy

For the STORM immunofluorescence, cells were fixed for 10 min with 4% paraformaldehyde and quenched with 10 mM ammonium chloride in 35 mm glass-bottomed dishes (81158, Ibidi). Cells were then permeabilized with 0.1% Triton X-100 in PBS for 7 min. The samples were then washed with PBS and blocked with PBS/2.5% goat serum/0.2% Tween 20 for 10 min, followed by 10 min of blocking with PBS/0.4% fish skin gelatin/0.2% Tween 20. Cells then were incubated with the primary antibody for 1 h at room temperature. Samples were washed 5 times with PBS/0.2% Tween 20 (2 times for 5 min and 3 times for 10 min) followed by 10 min of blocking with PBS/0.4% fish skin gelatin/0.2% Tween 20 and 10 min with PBS/2.5% goat serum/0.2% Tween 20. Secondary antibody (Alexa Fluor 488 and 647) diluted in the blocking solution was then added for 45 min, washed 5 times with PBS/0.2% Tween as before, and then washed twice with PBS to remove any detergent. Samples were imaged in OxEA buffer (Nahidiazar et al., 2016) for 10,000 frames at 100 Hz for each channel starting with the longer wavelength fluorophore first. The minimal light intensity to sufficiently send the fluorophores to the dark state was used to limit bleaching (Diekmann et al., 2020). Reconstruction of STORM data was performed using Nikon NIS-Elements AR software.

### FRET acquisition and processing

To monitor changes in Cdc42 activity in live cells we used a previously characterized dimerization-optimized reporter for activation (DORA) single chain Cdc42 biosensor (generous gift from Yi Wu, UConn Health, Farmington, CT) (Reinhard et al., 2017). Experiments were carried out as previously described for a similar Rac sensor (Baker et al., 2020; Cooke et al., 2021). Briefly, SUM159 cells stably expressing low levels of the FRET-sensor together with a marker for invadopodia (mCherry-cortactin), were treated with PDBu to induce invadopodia. We used 1:100 Oxyfluor reagent (Oxyrase Inc.) and 10 mM DL-lactate (Sigma-Aldrich) to reduce oxygen free radicals. Cells were imaged every 15 s with 2 x 2 binning and 16-bit depth, using a Nikon Eclipse Ti2 microscope equipped for FRET imaging (see Microscopes). Raw images were processed in batch using a custom designed macro in ImageJ, which included corrections to account for background and bleaching, and a median filter with a 1-pixel radius was applied to reduce noise artifacts. The FRET ratio was calculated, and a 16-color LUT was applied to allow for the visualization of Cdc42 activation.

### Microscopes

For STORM, TIRF, and live imaging experiments, a Nikon Eclipse Ti2 microscope was used equipped with a Tokai Hit STX stage top incubator, Apo TIRF 60x NA 1.49 oil immersion and SR HP Apo TIRF 100x NA 1.49 oil immersion objectives, Hamamatsu ORCA-Flash 4.0 camera, Princeton Instruments ProEM back-illuminated EMCCD camera, Lumencor SPECTRA X solid-state light source, Agilent laser unit (405-, 488-, 561-, and 647-nm), and NIS-Elements AR software. The microscope is equipped with a dichroic splitter to facilitate simultaneous acquisition of Cerulean3 and Venus emissions.

Confocal data was collected using a Leica Stellaris 5 laser scanning confocal equipped with an HC PL APO 63x/1.40 OIL CS2 objective and LAS X software, or an Andor Dragonfly 200 spinning disk confocal on a Leica DMI8 platform equipped with an HC PL FLUOTAR L 20X air objective, HC PL APO 63x/1.40 oil objective, Zyla4 camera, and Fusion software.

Brightfield images were acquired on an Olympus IX81 inverted microscope equipped with a UPlanFL N 10×/0.30 Ph1 air objective, Hamamatsu ORCA-Flash 4.0 camera, and cellSens software.

### Matrix degradation assay

The matrix degradation assay was modified from the Weed lab protocol (Martin et al., 2012). In brief, glass coverslips were coated with polylysine (50 µg/mL in water) and incubated for 20 minutes at room temperature and washed 3x with PBS. The coverslips were then fixed with ice-cold 0.5% glutaraldehyde in PBS and incubated on ice for 15 minutes and washed 3x with ice-cold PBS. The last wash was completely removed, and the coverslips were coated with a 37 °C solution consisting of 6 parts 1% (w/w) stock gelatin/sucrose solution in PBS, 1 part of Oregon Green 488-conjugated gelatin (Thermo Fisher Scientific), and 2 parts collagen IV (500 μg/mL stock). The excess matrix solution was aspirated off and incubated for 10 min at room temperature in the dark. The coverslips were then washed 3x with PBS and quenched with NaBH4 (5 mg/ml in PBS) for 15 min. The coverslips were washed 3x with PBS and sterilized with 70% ethanol for 30 min at room temperature. Finally, the coverslips were washed 3x with sterile PBS and stored at 4 °C for up to two weeks before use.

For the assay, SUM159 cells were plated on the gelatin coverslips and treated with PDBu for 24 hours. The coverslips were then fixed, stained with AlexaFluor-594 Phalloidin, and mounted using the described immunofluorescence protocol. For each condition, 20 random fields-of-view (FOV) were acquired via spinning disk confocal. Degradation was quantified by dividing the area of degradation for each image by the total cell area and normalizing the results to the CTRL cells.

### Invasion assays

The spheroid invasion assay was performed using the previously described method (Vinci et al., 2015). In brief, 4,000 cells were transferred into ultra-low attachment (ULA) 96-well round bottom plate in 200 μL of media and grown for 4 days. Once the spheroids had matured, 100 μL of media was removed and 100 μL of Matrigel (354230, Corning) was added and allowed to polymerase at 37 °C for 1 hr. After polymerization, 100 μL of media were added on top of the Matrigel. The spheroids were imaged every 24 hr for 72 hr. For measuring spheroid proliferation, the same process was followed except the Matrigel was omitted.

The inverse invasion assay was modified from previous methods (Goswami et al., 2005; Scott et al., 2011). SUM159 cells stably expressing mNeon-CAAX were plated in a CELLview glass bottom 10-well slide (Greiner Bio-One) to achieve 100% confluency overnight. The cells were then washed 3 times with PBS to remove growth factors and media. 100 μL of 50% Matrigel in serum free media was added to each of the wells. The cells were then incubated at 37 °C for 30 min to polymerize the matrix. 100 μL serum containing media was added to each well. The cells were then incubated overnight and at least 3 random regions per well were imaged with spinning disk confocal equipped with a long working distance 20X objective.

### Radial intensity analysis

Invadopodia radial intensity measurements were performed in ImageJ using a custom macro and the calculations were performed with RStudio. Briefly, a 3 μm line ROI was drawn across the center of a single invadopodia based on the F-actin channel. Then the pixel intensity was measured across the ROI after which the ROI was rotated 3 degrees. This was repeated until the ROI completed a complete revolution and was then repeated for each channel of the image. For each invadopodia measured, the intensity values were averaged together to calculate the average intensity radiating out from the center of the invadopodia. The intensity measurements can then be plotted directly or used to calculate the distance of the peak signal from the center of each invadopodia or the relative signal intensity for each invadopodia by dividing the average signal intensity within 1 μm from the center of the invadopodia by the signal 1.5 μm from the invadopodia center.

### Invadopodia dynamics analysis

The invadopodia dynamics measurements were performed in ImageJ using a custom macro and the calculations were performed with RStudio. Briefly, the center of a random, single invadopodia that forms and disassembles within the duration of the video was identified for each frame. Then, for each frame, a 1 μm diameter circle was drawn around the invadopodia and a 1 μm band was drawn around the circle. The average intensity within the circle was then divided by the average intensity within the band to calculate the relative signal intensity at the invadopodia for each frame. The quantification of the dynamics was performed as previously described for cell ruffle dynamics (Kreider-Letterman et al., 2022b).

### Statistical analysis

Values calculated from at least three independent experiments were compared by an unpaired student’s t-test using GraphPad Prism or RStudio unless otherwise indicated. Otherwise, a one-way ANOVA analysis followed by a Tukey’s test was performed. Technical replicates are defined in this study as individual cells or invadopodia while biological replicates are defined as the averages of technical replicates or cell populations from independently prepared samples. Biological replicates were assumed to have normal distribution (Fay and Gerow, 2013). P<0.05 was considered statistically significant. Error bars represent the S.E.M. unless otherwise indicated.

## Acknowledgements

The authors would like to thank Carol Otey, Harvey T. McMahon, Jaap van Buul, and Lou Hodgson for providing reagents. We would also like to thank Jaap van Buul, Jose J. Bravo-Cordero, Marcelo Kazanietz, as well as the members of the Garcia-Mata and Goedhart lab for critical comments and valuable discussion. This work was supported by a Nederlandse Organisatie voor Wetenschappelijk Onderzoek ALW-OPEN grant (ALWOP.306) (to E.K.M.) and 1R15CA199101, R03CA234693, and R01-GM136826-01 grants from the NIH (to R.G.-M.).

## Supplementary figure legends

**Figure S1.**
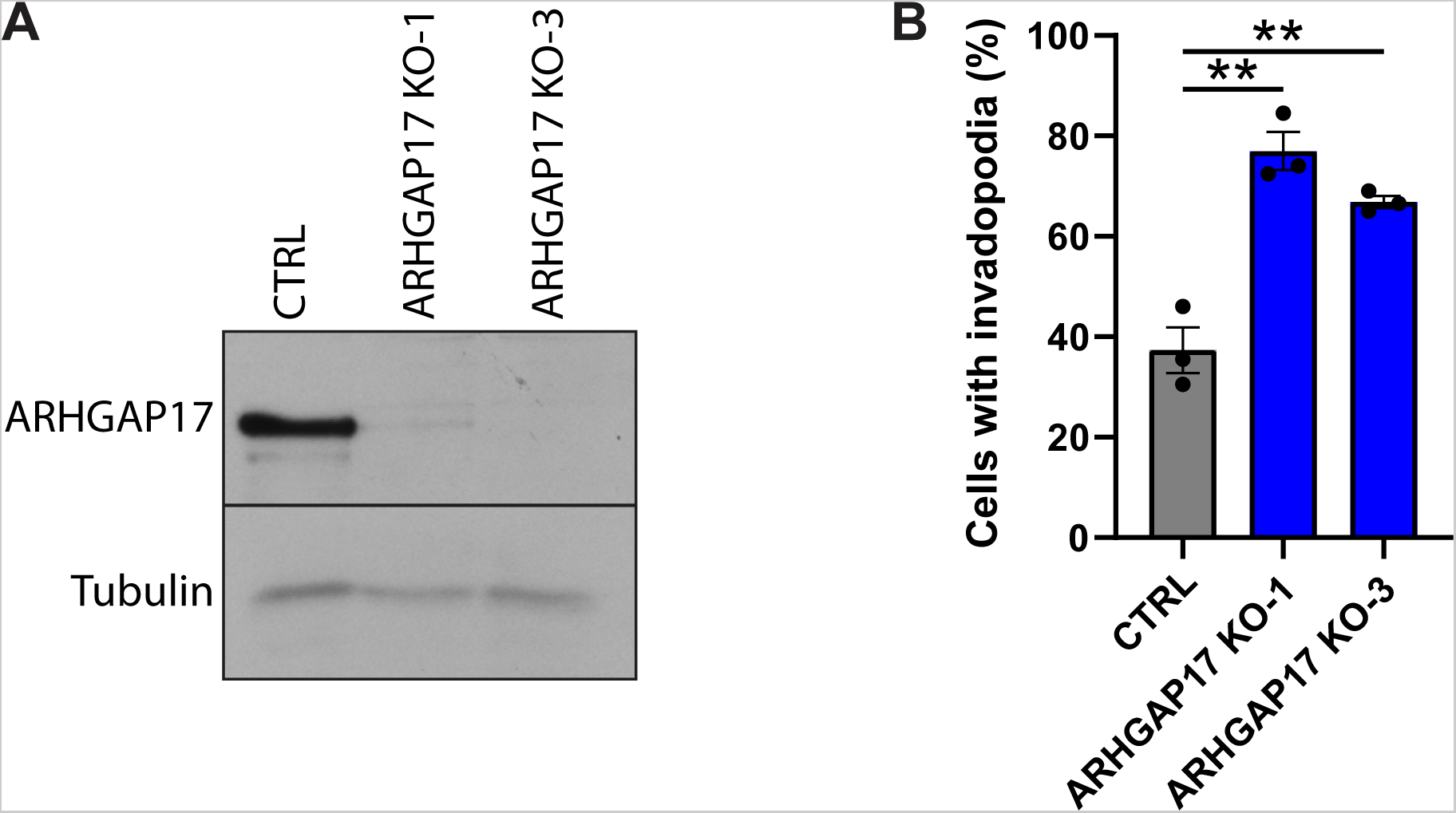
ARHGAP17 KO validation. (**A**) Representative western blot showing loss of ARHGAP17 protein in SUM159 cells using two different gRNA targets. (**B**) Percentage of cells with invadopodia after treatment with PDBu for 30 minutes. Results are expressed as mean ± S.E.M. of 3 independent experiments in which at least 200 cells per condition per experiment were counted. **, p<0.01.

**Figure S2.**
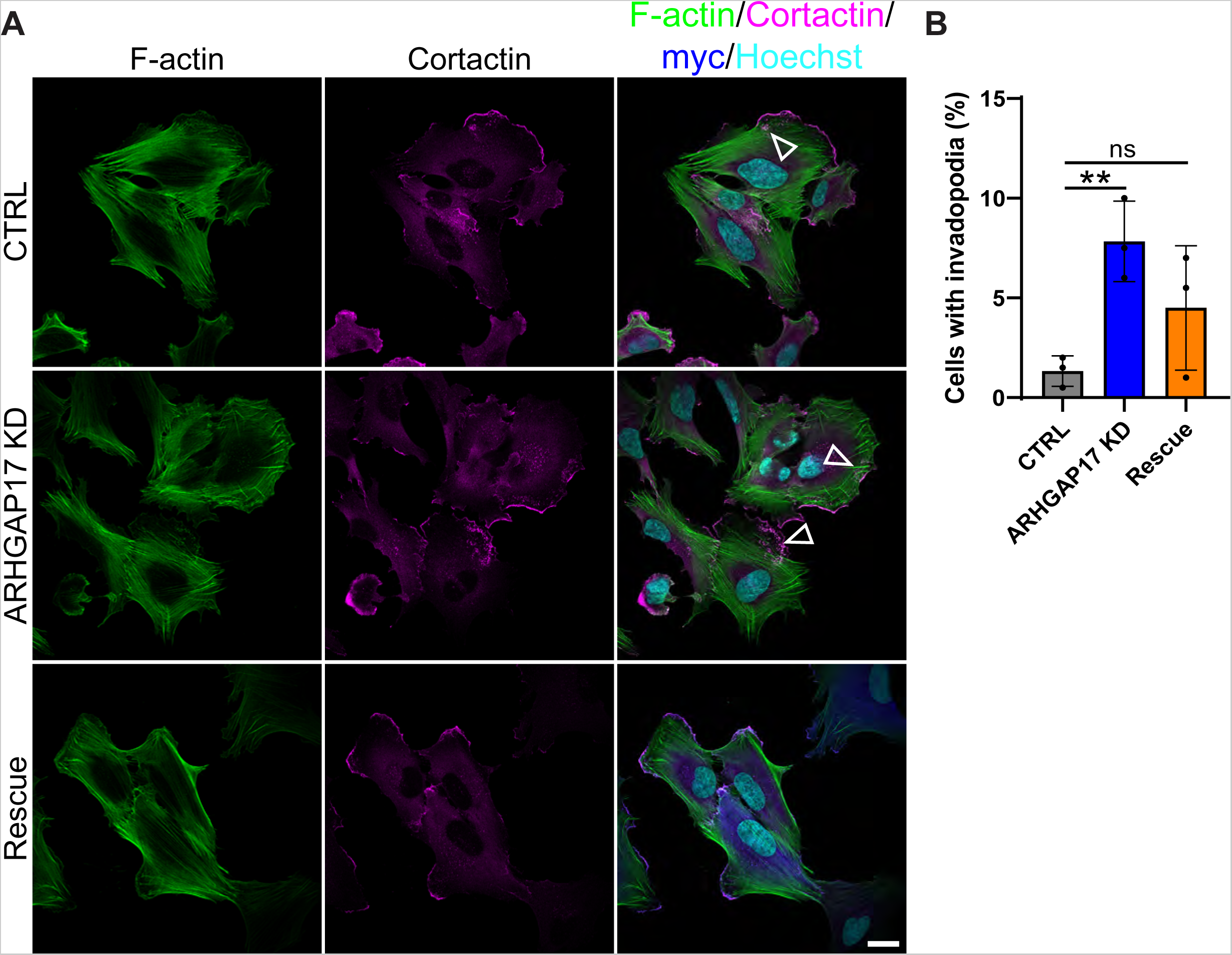
ARHGAP17 KD increases the formation of spontaneous invadopodia. (**A**) Representative micrographs of ARHGAP17 knockdown and Rescue with shRNA resistant myc-ARHGAP17 in SUM159 cells. F-actin and cortactin staining were used as markers for invadopodia while myc staining was used to visualize myc-ARHGAP17. Arrowheads highlight invadopodia clusters (scale bar, 20 μm). (**B**) Percentage of cells with invadopodia after treatment with PDBu for 30 minutes. Results are expressed as mean ± S.E.M. of 3 independent experiments in which at least 200 cells per condition per experiment were counted. **, p<0.01; ns, not significant.

**Figure S3.**
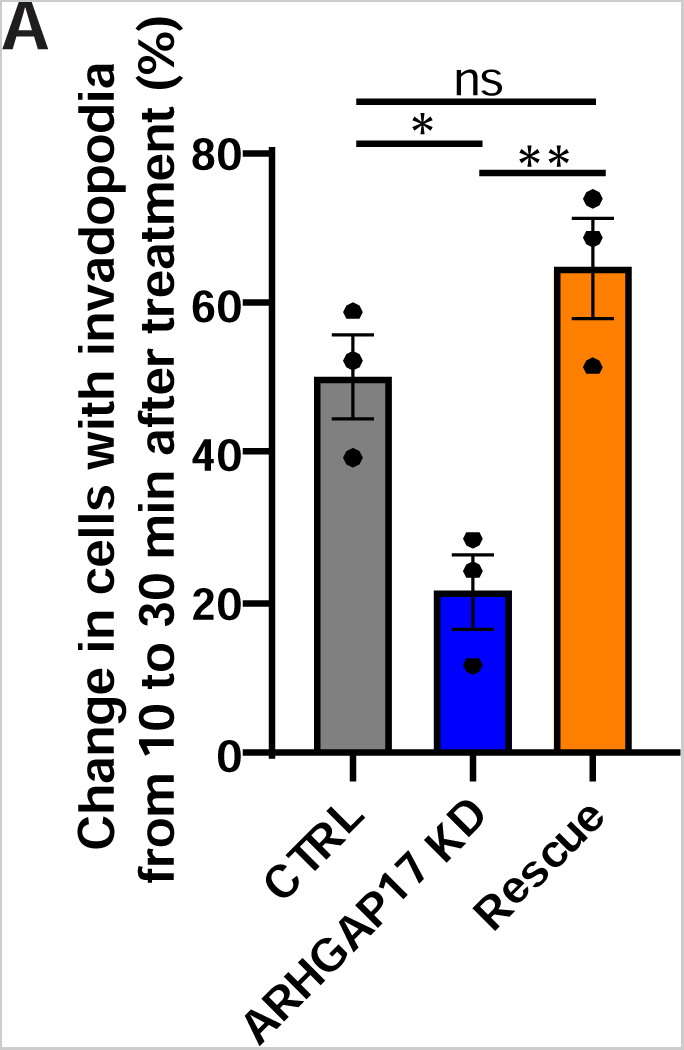
ARHGAP17 KD reduces invadopodia turnover. (**A**) Quantification of change in percentage of cells forming invadopodia between peak invadopodia formation 10 minutes after PDBu treatment and equilibrium at 30 minutes after PDBu treatment. Results are expressed as mean ± S.E.M. of 3 independent experiments in which at least 200 cells per condition per experiment were counted. *, p<0.05; **, p<0.01; ns, not significant.

**Figure S4.**
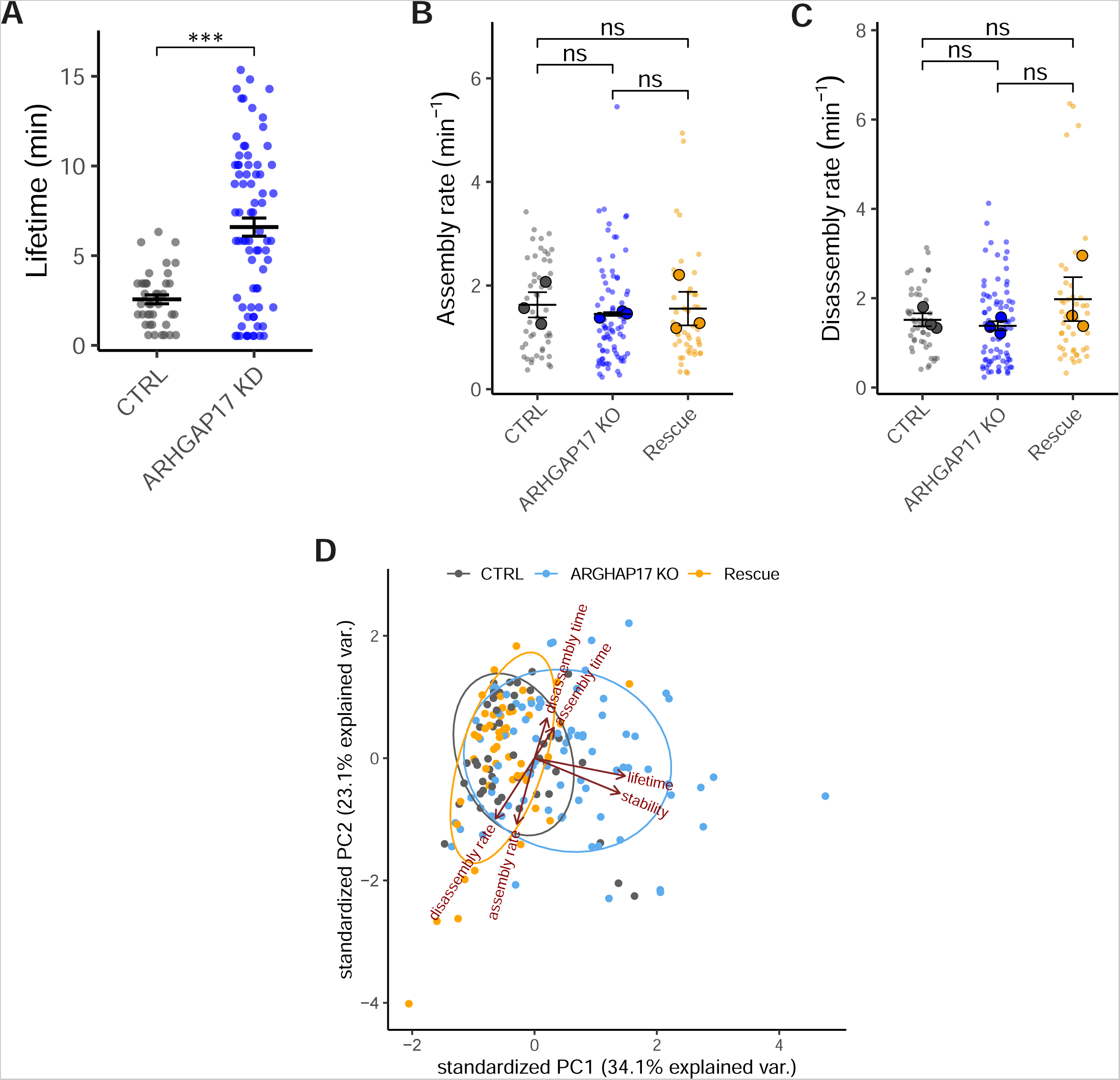
ARHGAP17 regulates invadopodia lifetime and stability. (**A**) Quantification of invadopodia lifetime in untreated SUM159 cells. Results are expressed as mean ± S.E.M of at least 40 invadopodia per condition. ***, p<0.001. (**B and C**) Quantification of invadopodia assembly and disassembly rate. Results are expressed as mean ± S.E.M. of 3 independent experiments. Large circles represent biological replicates, small circles represent technical replicates. ns, not significant. (**D**) Principal component analysis of invadopodia dynamics. The ellipses represent 95% confidence.

**Figure S5.**
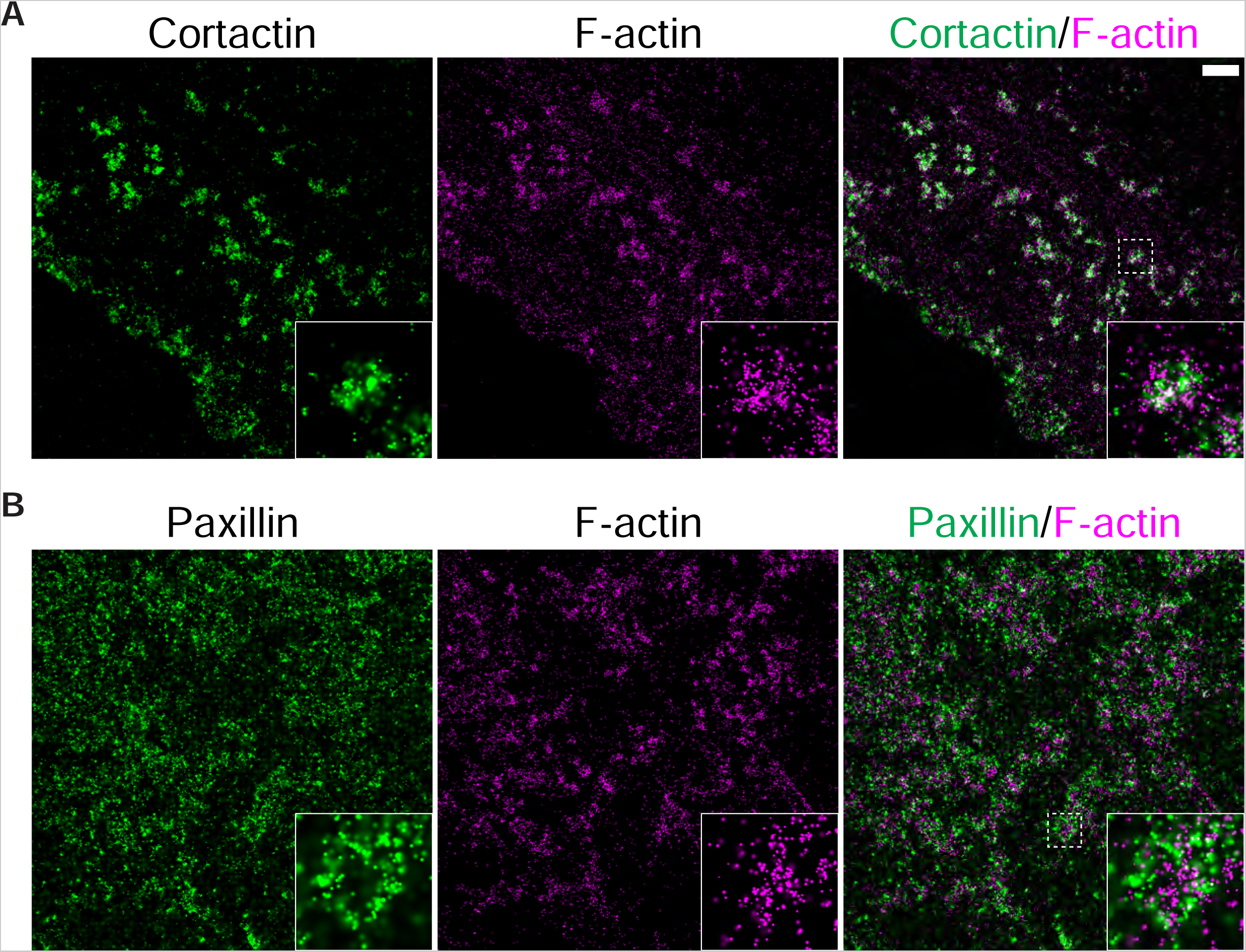
Paxillin and cortactin localization to invadopodia. (**A and B**) STORM reconstructions of SUM159 cells treated with PDBu showing cortactin and paxillin localization to invadopodia as defined by F-actin staining (scale bar, 1 μm).

**Figure S6.**
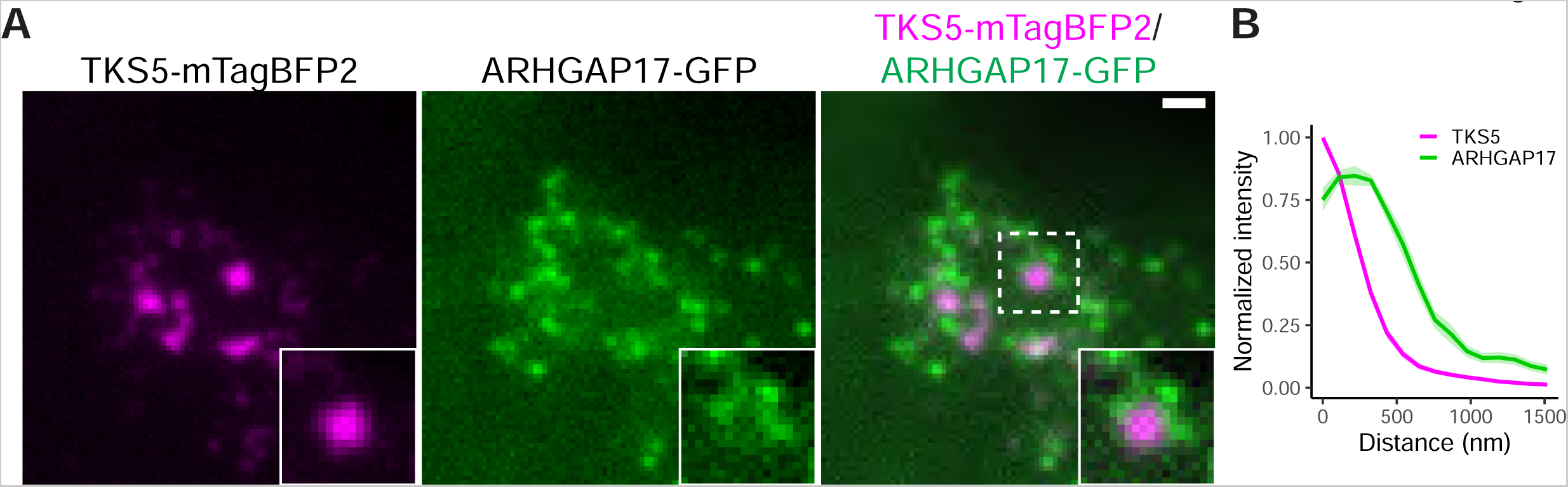
ARHGAP17-GFP localization at invadopodia in untreated cells. (**A**) Example micrograph of a SUM159 cell invadopodia cluster expressing TKS5-mTagBFP2 and ARHGAP17-GFP in an ARHGAP17 KO background (scale bar, 1 μm). (**B**) Radial intensity analysis of TKS5-mTagBFP2 and ARHGAP17-GFP at invadopodia. Lines are expressed as mean ± S.E.M. of 3 independent experiments in which at least 10 invadopodia per condition per experiment were measured.

**Figure S7.**
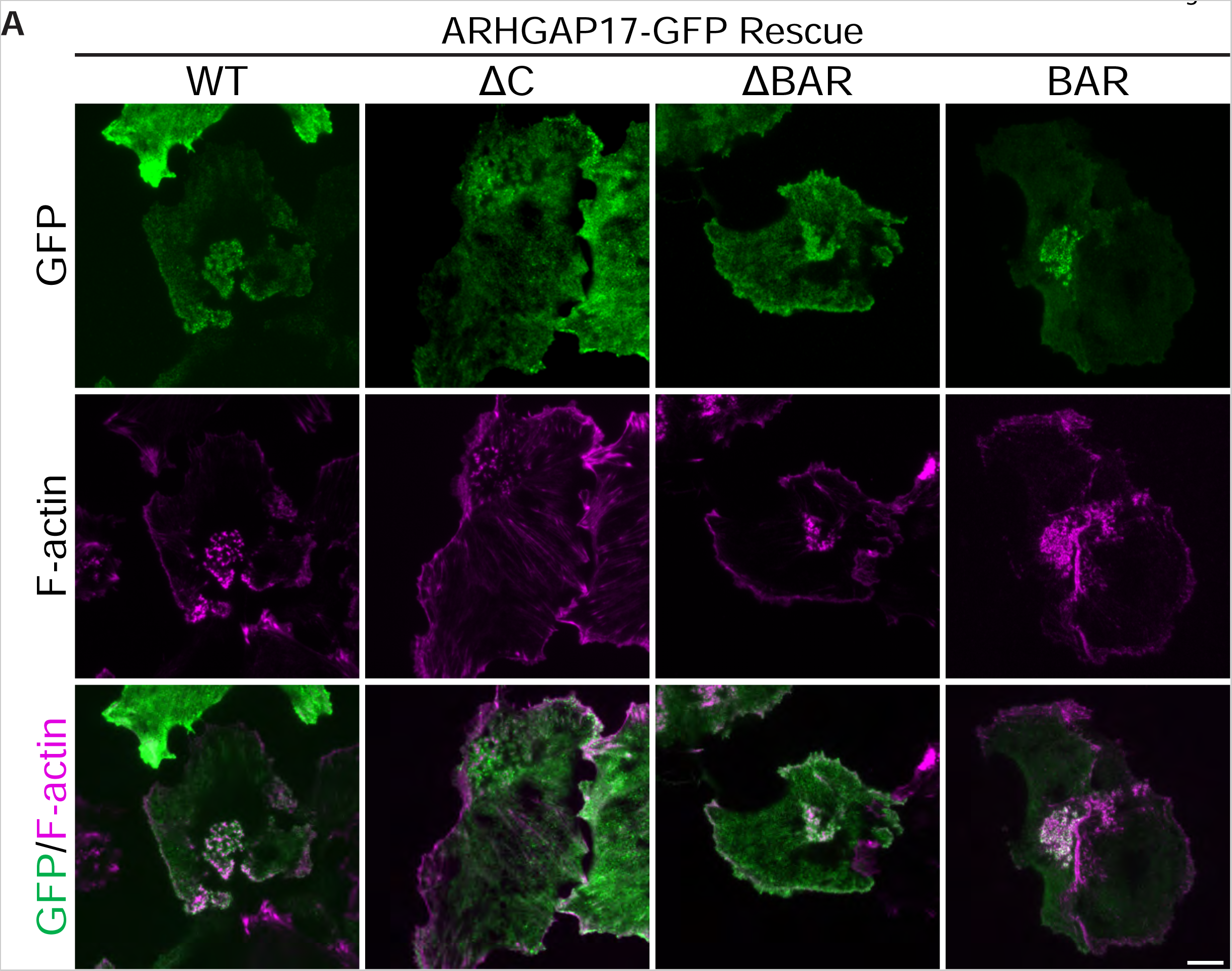
Localization of ARHGAP17 domain deletion mutants. (**A**) Representative TIRF micrographs of SUM159 cells expressing ARHGAP17-GFP domain deletion mutants in an ARHGAP17 KO background. Cells were treated with PDBu for 30 min and stained for F-actin as a marker for invadopodia. Arrowheads highlight invadopodia clusters (scale bar for both panels, 10 μm).

**Figure S8.**
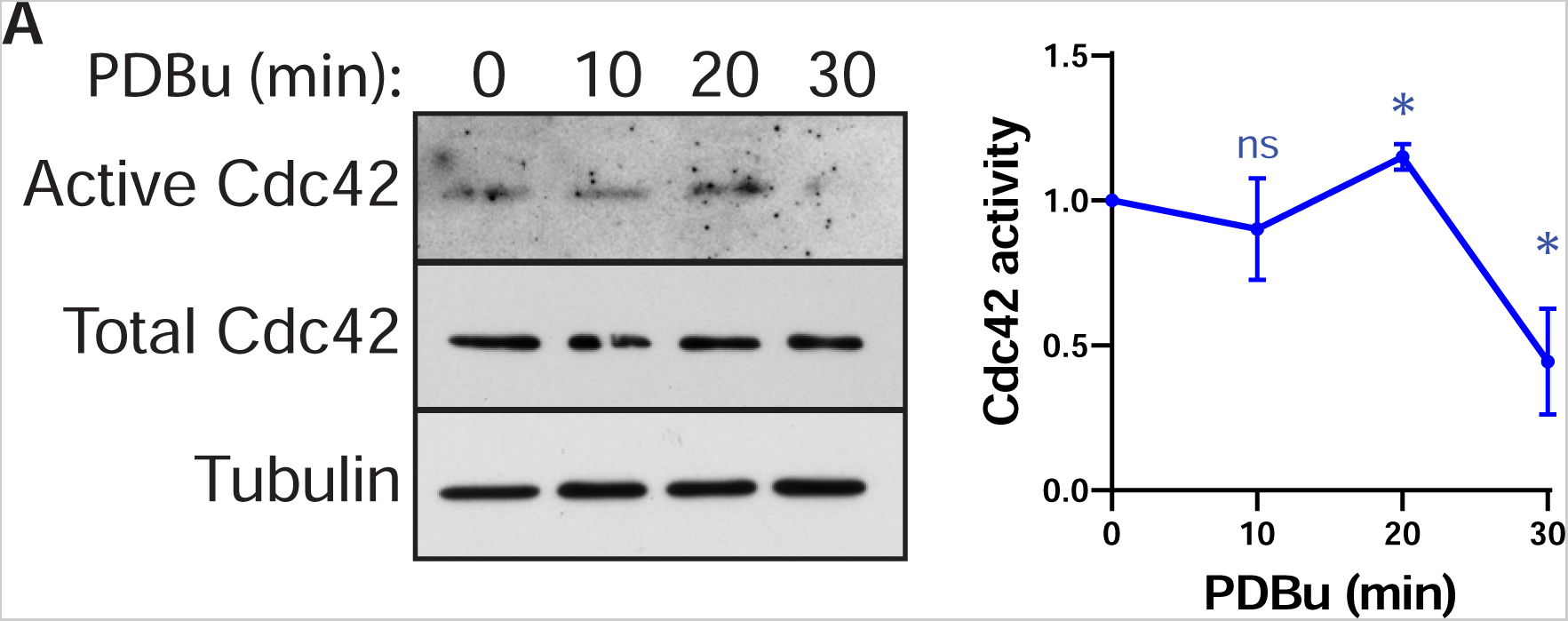
Cdc42 activity in ARHGAP17 KD cells. (**A**) PBD pulldown of active Cdc42 in ARHGAP17 KD SUM159 cells after treatment with PDBu at the indicated time points. Results are expressed as mean ± S.E.M. of 3 independent experiments. *, p<0.05; ns, not significant.

**Figure S9.**
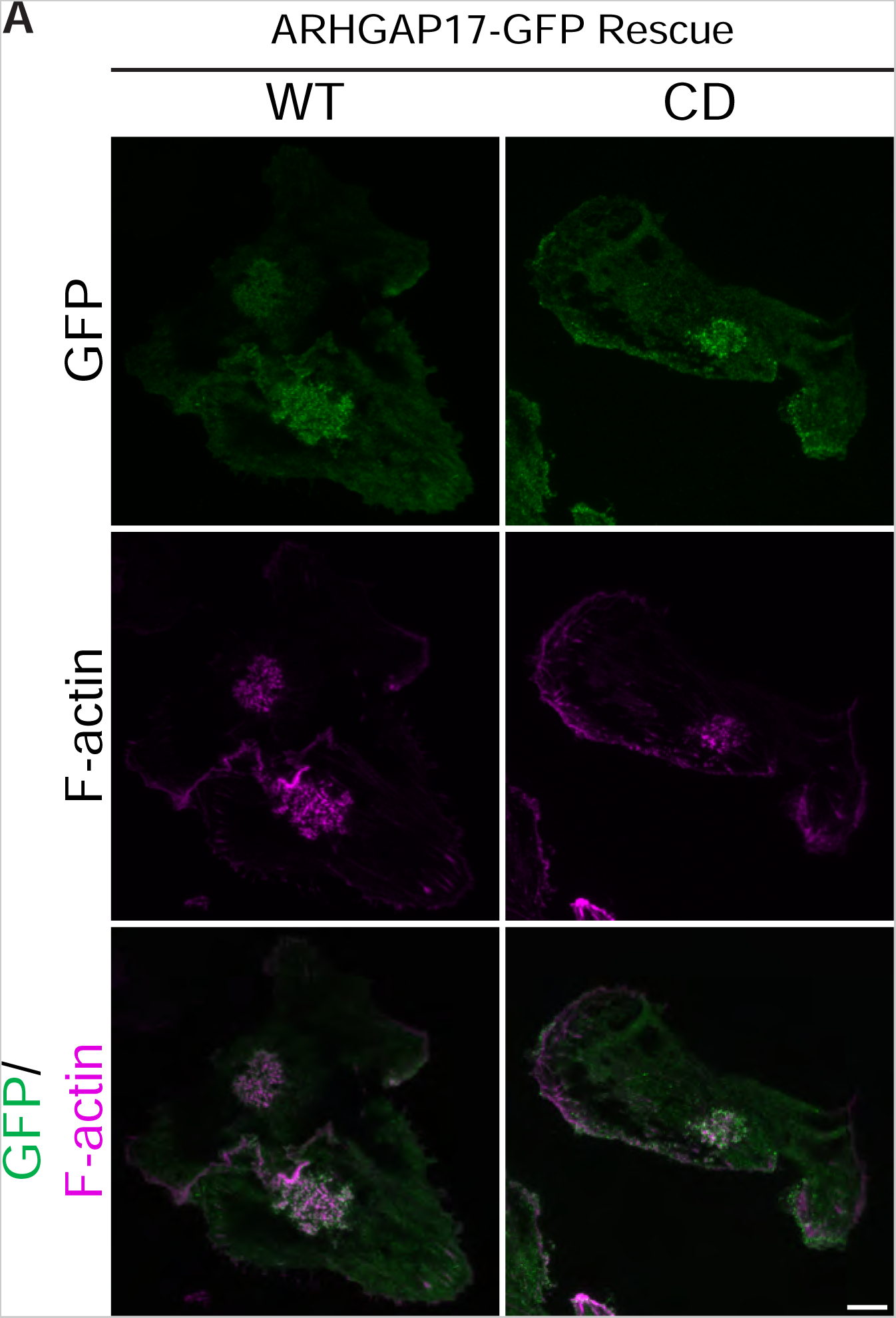
Localization of ARHGAP17 CD. (**A**) Representative TIRF micrographs of SUM159 cells expressing FL-ARHGAP17-GFP and CD-ARHGAP17-GFP in an ARHGAP17 KO background. Cells were treated with PDBu for 30 min and stained for F-actin as a marker for invadopodia. Arrowheads highlight invadopodia clusters (scale bar for both panels, 10 μm).

**Figure S10.**
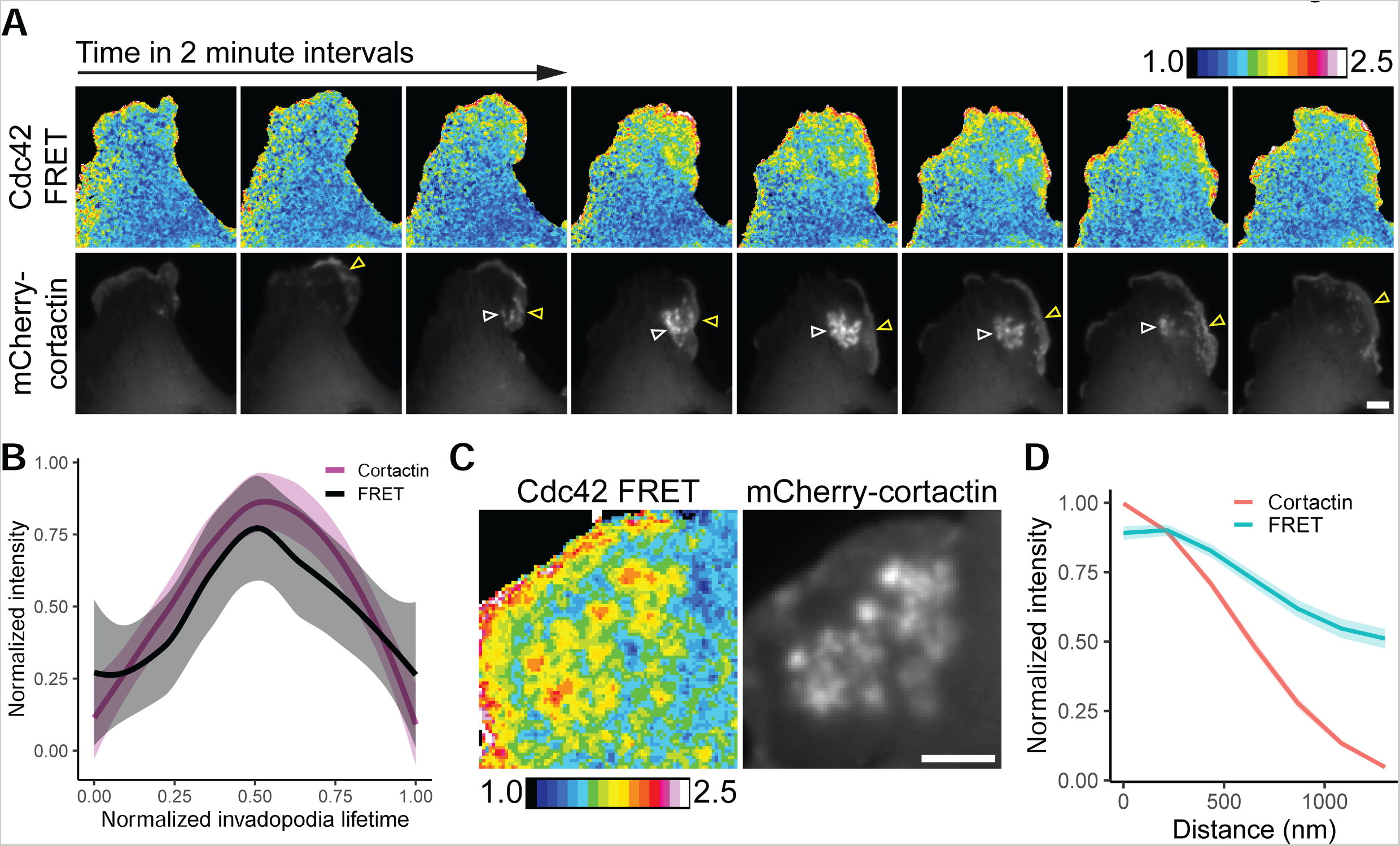
Cdc42 FRET biosensor activity at invadopodia. (**A**) Representative micrographs of a SUM159 cell expressing the single-chain Cdc42 FRET biosensor and mCherry-cortactin as a marker for invadopodia after PDB treatment (scale bar, 5 μm). The color calibration bar indicates the relative FRET ratio. White arrowheads indicate invadopodia cluster, yellow arrowheads indicate lamellipodia protrusion. (**B**) Quantification of FRET ratio and mCherry-cortactin intensity at individual invadopodia. Intensity and time have been rescaled between values of 0 and 1 for each invadopodia to allow for comparison. Lines are loess smoothing and 95% confidence from 5 invadopodia. (**C**) Example frame highlighting Cdc42 FRET at invadopodia in a SUM159 cell as marked by mCherry-cortactin (scale bar, 5 μm). (**D**) Radial intensity analysis of FRET ratio and mCherry-cortactin intensity at invadopodia. Lines are expressed as mean ± S.E.M. of 40 invadopodia.

**Figure S11.**
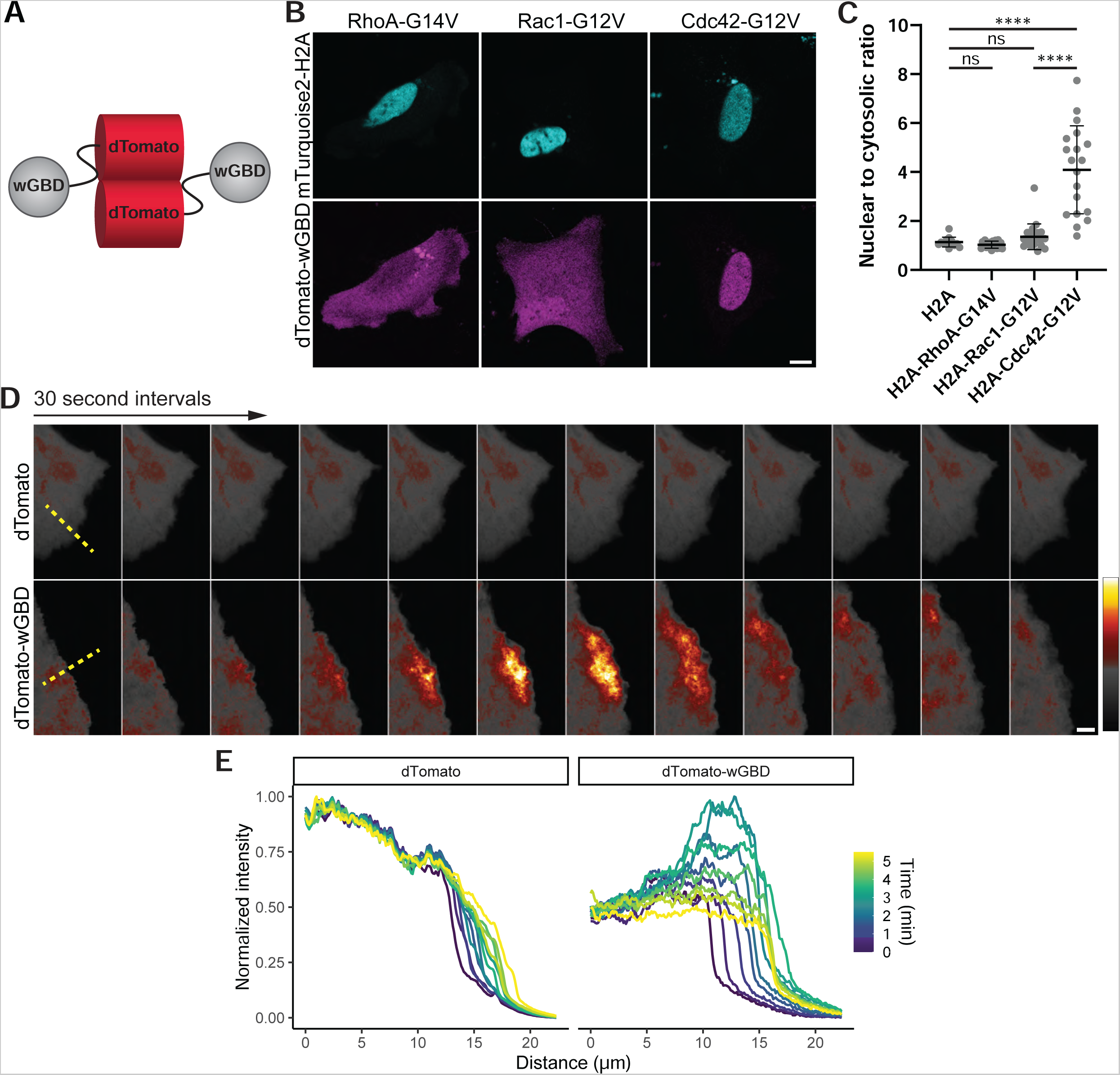
Validation of the Cdc42 biosensor specificity. (**A**) Design of the Cdc42 activity biosensor used in this study, composed of the GTPase-binding domain of WASP (wGBD) tagged with dTomato. (**B**) Representative micrographs of SUM159 cells showing colocalization of the dTomato-wGBD sensor with different mTurquoise2-H2A-RhoGTPase-ΔCAAX constructs (RhoA-G14V, Rac1-G12V, or Cdc42-G12V) (scale bar, 10 μm). (**C**) Quantification of the average intensity of dTomato-wGBD signal within the nucleus ratioed to the average intensity in the cytosol. Results are mean ± S.D. of at least 14 cells per condition from 3 independent experiments. ****, p<0.0001; ns, not significant. (**D**) Representative TIRF micrographs of lamellipodia protrusions from SUM159 cells expressing either dTomato as a control for cytoplasmic localization or dTomato-wGBD (scale bar, 5 μm). (**E**) Quantification of panel (**D**) in which a 10-pixel wide line was drawn perpendicular to the cell edge (yellow dashed line in panel D) and the intensity was measured for each frame. Distance 0 starts 10-15 μm within the cell and the protrusion can be visualized by a change in distance at intensity 0.25 over time. The increase in peak dTomato-wGBD intensity over time indicates an increase in Cdc42 activity localized to the leading edge.

**Figure S12.**
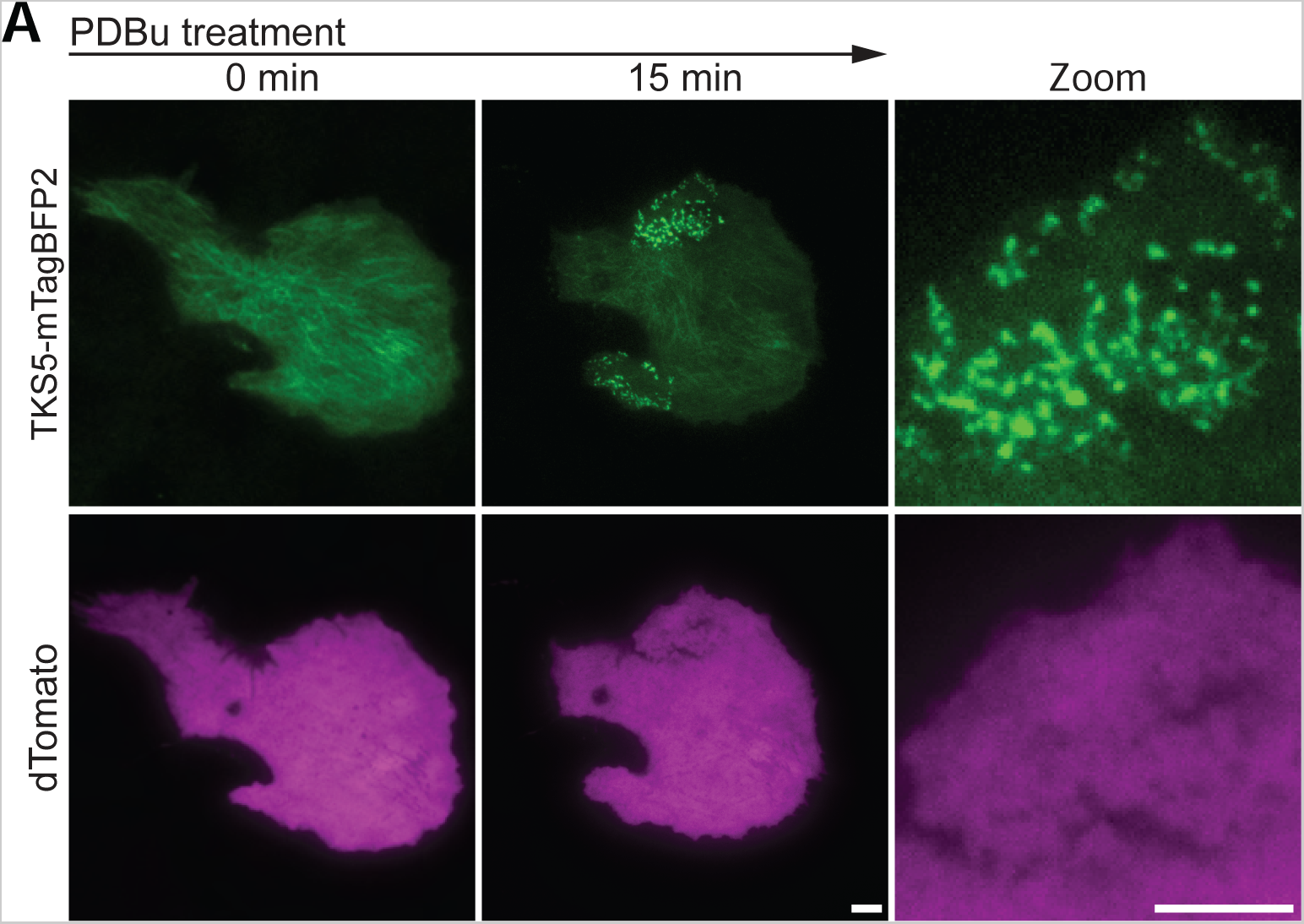
Control for dTomato-wGBD invadopodia localization. (**A**) Representative TIRF micrographs of a SUM159 cell expressing TKS5-mTagBFP2 as a marker for invadopodia and dTomato as a localization control for the dTomato-wGBD biosensor forming an invadopodia cluster after PDBu treatment (scale bar for both panels, 5 μm).

## Video figure legends

Video 1. SUM159 cell forming invadopodia after PDBu treatment

Live TIRF imaging of a SUM159 cell expressing TKS5-mTagBFP2 and treated with PDBu (scale bar, 10 μm).

Video 2. SUM159 cell inverse invasion assay

Live spinning-disk z-projection of SUM159 cells expressing mScarlet-CAAX invading into Matrigel (scale bar, 10 μm).

Video 3. ARHGAP17 localizes to invadopodia

Live TIRF imaging of a SUM159 cell expressing TKS5-mTagBFP2 and ARHGAP17-GFP (expressed in ARHGAP17 KO background), treated with PDBu (scale bar, 10 μm).

Video 4. dTomato-wGBD sensor localization to membrane protrusions

Live TIRF imaging of membrane protrusions from SUM159 cells expressing either dTomato as a control for cytoplasmic localization or dTomato-wGBD (scale bar, 5 μm).

Video 5. dTomato-wGBD sensor localizes to invadopodia.

Live TIRF imaging of a SUM159 cell expressing TKS5-mTagBFP2 and dTomato-wGBD, treated with PDBu (scale bar, 10 μm).

Video 6. dTomato does not localize to invadopodia.

Live TIRF imaging of a SUM159 cell expressing TKS5-mTagBFP2 and dTomato, treated with PDBu (scale bar, 10 μm).

Video 7. Localization of ARHGAP17 and Cdc42 activity at an individual invadopodia.

Live TIRF of an individual invadopodia from a SUM159 cells expressing TKS5-mTagBFP2, ARHGAP17-GFP (expressed in ARHGAP17 KO background), and dTomato-wGBD, treated with PDBu (scale bar, 1 μm).

Video 8. Localization of CIP4 and Cdc42 activity at an individual invadopodia.

Live TIRF of an individual invadopodia from a SUM159 cells expressing TKS5-mTagBFP2, GFP-CIP4, and dTomato-wGBD, treated with PDBu (scale bar, 1 μm).

## References

1. Adikusuma, F., C. Pfitzner, and P.Q. Thomas. 2017. Versatile single-step-assembly CRISPR/Cas9 vectors for dual gRNA expression. PLoS One. 12:e0187236.

2. Al Haddad, M., R. El-Rif, S. Hanna, L. Jaafar, R. Dennaoui, S. Abdellatef, V. Miskolci, D. Cox, L. Hodgson, and M. El-Sibai. 2020. Differential regulation of rho GTPases during lung adenocarcinoma migration and invasion reveals a novel role of the tumor suppressor StarD13 in invadopodia regulation. Cell Commun Signal. 18:144.

3. Artym, V.V., Y. Zhang, F. Seillier-Moiseiwitsch, K.M. Yamada, and S.C. Mueller. 2006. Dynamic interactions of cortactin and membrane type 1 matrix metalloproteinase at invadopodia: defining the stages of invadopodia formation and function. Cancer Res. 66:3034–3043.

4. Aspenstrom, P. 2018. BAR Domain Proteins Regulate Rho GTPase Signaling. Adv Exp Med Biol.

5. Ayala, I., G. Giacchetti, G. Caldieri, F. Attanasio, S. Mariggio, S. Tete, R. Polishchuk, V. Castronovo, and R. Buccione. 2009. Faciogenital dysplasia protein Fgd1 regulates invadopodia biogenesis and extracellular matrix degradation and is up-regulated in prostate and breast cancer. Cancer Res. 69:747–752.

6. Badowski, C., G. Pawlak, A. Grichine, A. Chabadel, C. Oddou, P. Jurdic, M. Pfaff, C. Albigès-Rizo, and M.R. Block. 2008. Paxillin phosphorylation controls invadopodia/podosomes spatiotemporal organization. Mol Biol Cell. 19:633–645.

7. Baker, M.J., M. Cooke, G. Kreider-Letterman, R. Garcia-Mata, P.A. Janmey, and M.G. Kazanietz. 2020. Evaluation of active Rac1 levels in cancer cells: a case of misleading conclusions from immunofluorescence analysis. Journal of Biological Chemistry.

8. Benink, H.A., and W.M. Bement. 2005. Concentric zones of active RhoA and Cdc42 around single cell wounds. J Cell Biol. 168:429–439.

9. Berginski, M.E., E.A. Vitriol, K.M. Hahn, and S.M. Gomez. 2011. High-resolution quantification of focal adhesion spatiotemporal dynamics in living cells. PLoS One. 6:e22025.

10. Branch, K.M., D. Hoshino, and A.M. Weaver. 2012. Adhesion rings surround invadopodia and promote maturation. Biol Open. 1:711–722.

11. Bravo-Cordero, J.J., M. Oser, X. Chen, R. Eddy, L. Hodgson, and J. Condeelis. 2011. A novel spatiotemporal RhoC activation pathway locally regulates cofilin activity at invadopodia. Curr Biol. 21:635–644.

12. Bu, W., K.B. Lim, Y.H. Yu, A.M. Chou, T. Sudhaharan, and S. Ahmed. 2010. Cdc42 interaction with N-WASP and Toca-1 regulates membrane tubulation, vesicle formation and vesicle motility: implications for endocytosis. PLoS One. 5:e12153.

13. Chan Wah Hak, L., S. Khan, I. Di Meglio, A.L. Law, S. Lucken-Ardjomande Hasler, L.M. Quintaneiro, A.P.A. Ferreira, M. Krause, H.T. McMahon, and E. Boucrot. 2018. FBP17 and CIP4 recruit SHIP2 and lamellipodin to prime the plasma membrane for fast endophilin-mediated endocytosis. Nat Cell Biol. 20:1023–1031.

14. Chander, H., P. Truesdell, J. Meens, and A.W. Craig. 2013. Transducer of Cdc42-dependent actin assembly promotes breast cancer invasion and metastasis. Oncogene. 32:3080–3090.

15. Cooke, M., G. Kreider-Letterman, M.J. Baker, S. Zhang, N.T. Sullivan, E. Eruslanov, M.C. Abba, S.M. Goicoechea, R. García-Mata, and M.G. Kazanietz. 2021. FARP1, ARHGEF39, and TIAM2 are essential receptor tyrosine kinase effectors for Rac1-dependent cell motility in human lung adenocarcinoma. Cell Rep. 37:109905.

16. Daubon, T., R. Buccione, and E. Génot. 2011. The Aarskog-Scott syndrome protein Fgd1 regulates podosome formation and extracellular matrix remodeling in transforming growth factor β-stimulated aortic endothelial cells. Mol Cell Biol. 31:4430–4441.

17. Desmarais, V., H. Yamaguchi, M. Oser, L. Soon, G. Mouneimne, C. Sarmiento, R. Eddy, and J. Condeelis. 2009. N-WASP and cortactin are involved in invadopodium-dependent chemotaxis to EGF in breast tumor cells. Cell Motil Cytoskeleton. 66:303–316.

18. Di Martino, J., L. Paysan, C. Gest, V. Lagrée, A. Juin, F. Saltel, and V. Moreau. 2014. Cdc42 and Tks5: a minimal and universal molecular signature for functional invadosomes. Cell Adh Migr. 8:280–292.

19. Diekmann, R., M. Kahnwald, A. Schoenit, J. Deschamps, U. Matti, and J. Ries. 2020. Optimizing imaging speed and excitation intensity for single-molecule localization microscopy. Nat Methods. 17:909–912.

20. Diring, J., S. Mouilleron, N.Q. McDonald, and R. Treisman. 2019. RPEL-family rhoGAPs link Rac/Cdc42 GTP loading to G-actin availability. Nat Cell Biol. 21:845–855.

21. Dombrosky-Ferlan, P., A. Grishin, R.J. Botelho, M. Sampson, L. Wang, W.A. Rudert, S. Grinstein, and S.J. Corey. 2003. Felic (CIP4b), a novel binding partner with the Src kinase Lyn and Cdc42, localizes to the phagocytic cup. Blood. 101:2804–2809.

22. Fay, D.S., and K. Gerow. 2013. A biologist’s guide to statistical thinking and analysis. WormBook:1–54.

23. García-Mata, R., K. Wennerberg, W.T. Arthur, N.K. Noren, S.M. Ellerbroek, and K. Burridge. 2006. Analysis of activated GAPs and GEFs in cell lysates. Methods Enzymol. 406:425–437.

24. Gligorijevic, B., A. Bergman, and J. Condeelis. 2014. Multiparametric classification links tumor microenvironments with tumor cell phenotype. PLoS Biol. 12:e1001995.

25. Goicoechea, S.M., B. Bednarski, R. Garcia-Mata, H. Prentice-Dunn, H.J. Kim, and C.A. Otey. 2009. Palladin contributes to invasive motility in human breast cancer cells. Oncogene. 28:587–598.

26. Goicoechea, S.M., R. García-Mata, J. Staub, A. Valdivia, L. Sharek, C.G. McCulloch, R.F. Hwang, R. Urrutia, J.J. Yeh, H.J. Kim, and C.A. Otey. 2014. Palladin promotes invasion of pancreatic cancer cells by enhancing invadopodia formation in cancer-associated fibroblasts. Oncogene. 33:1265–1273.

27. Goicoechea, S.M., A. Zinn, S.S. Awadia, K. Snyder, and R. Garcia-Mata. 2017. A RhoG-mediated signaling pathway that modulates invadopodia dynamics in breast cancer cells. J Cell Sci. 130:1064–1077.

28. Goswami, S., E. Sahai, J.B. Wyckoff, M. Cammer, D. Cox, F.J. Pixley, E.R. Stanley, J.E. Segall, and J.S. Condeelis. 2005. Macrophages promote the invasion of breast carcinoma cells via a colony-stimulating factor-1/epidermal growth factor paracrine loop. Cancer Res. 65:5278–5283.

29. Guo, Q., Y. Xiong, Y. Song, K. Hua, and S. Gao. 2019. ARHGAP17 suppresses tumor progression and up-regulates P21 and P27 expression via inhibiting PI3K/AKT signaling pathway in cervical cancer. Gene. 692:9–16.

30. Harada, A., B. Furuta, K. Takeuchi, M. Itakura, M. Takahashi, and M. Umeda. 2000. Nadrin, a novel neuron-specific GTPase-activating protein involved in regulated exocytosis. J Biol Chem. 275:36885–36891.

31. Hartig, S.M., S. Ishikura, R.S. Hicklen, Y. Feng, E.G. Blanchard, K.A. Voelker, C.S. Pichot, R.W. Grange, R.M. Raphael, A. Klip, and S.J. Corey. 2009. The F-BAR protein CIP4 promotes GLUT4 endocytosis through bidirectional interactions with N-WASp and Dynamin-2. J Cell Sci. 122:2283–2291.

32. Ho, H.Y., R. Rohatgi, A.M. Lebensohn, M. Le, J. Li, S.P. Gygi, and M.W. Kirschner. 2004. Toca-1 mediates Cdc42-dependent actin nucleation by activating the N-WASP-WIP complex. Cell. 118:203–216.

33. Hoshino, D., K.M. Branch, and A.M. Weaver. 2013. Signaling inputs to invadopodia and podosomes. J Cell Sci. 126:2979–2989.

34. Hoshino, D., J. Jourquin, S.W. Emmons, T. Miller, M. Goldgof, K. Costello, D.R. Tyson, B. Brown, Y. Lu, N.K. Prasad, B. Zhang, G.B. Mills, W.G. Yarbrough, V. Quaranta, M. Seiki, and A.M. Weaver. 2012. Network analysis of the focal adhesion to invadopodia transition identifies a PI3K-PKCα invasive signaling axis. Sci Signal. 5:ra66.

35. Hu, J., A. Mukhopadhyay, and A.W. Craig. 2011. Transducer of Cdc42-dependent actin assembly promotes epidermal growth factor-induced cell motility and invasiveness. J Biol Chem. 286:2261–2272.

36. Juin, A., C. Billottet, V. Moreau, O. Destaing, C. Albiges-Rizo, J. Rosenbaum, E. Genot, and F. Saltel. 2012. Physiological type I collagen organization induces the formation of a novel class of linear invadosomes. Mol Biol Cell. 23:297–309.

37. Juin, A., J. Di Martino, B. Leitinger, E. Henriet, A.S. Gary, L. Paysan, J. Bomo, G. Baffet, C. Gauthier-Rouvière, J. Rosenbaum, V. Moreau, and F. Saltel. 2014. Discoidin domain receptor 1 controls linear invadosome formation via a Cdc42-Tuba pathway. J Cell Biol. 207:517–533.

38. Kedziora, K.M., D. Leyton-Puig, E. Argenzio, A.J. Boumeester, B. van Butselaar, T. Yin, Y.I. Wu, F.N. van Leeuwen, M. Innocenti, K. Jalink, and W.H. Moolenaar. 2016. Rapid Remodeling of Invadosomes by Gi-coupled Receptors: DISSECTING THE ROLE OF Rho GTPases. J Biol Chem. 291:4323–4333.

39. Kim, S.H., Z. Li, and D.B. Sacks. 2000. E-cadherin-mediated cell-cell attachment activates Cdc42. J Biol Chem. 275:36999–37005.

40. Kiso, M., S. Tanaka, S. Saji, M. Toi, and F. Sato. 2018. Long isoform of VEGF stimulates cell migration of breast cancer by filopodia formation via NRP1/ARHGAP17/Cdc42 regulatory network. Int J Cancer.

41. Kreider-Letterman, G., N.M. Carr, and R. Garcia-Mata. 2022a. Fixing the GAP: The role of RhoGAPs in cancer. Eur J Cell Biol. 101:151209.

42. Kreider-Letterman, G., M. Cooke, S.M. Goicoechea, M.G. Kazanietz, and R. Garcia-Mata. 2022b. Quantification of ruffle area and dynamics in live or fixed lung adenocarcinoma cells. STAR Protocols. 3:101437.

43. Lawson, C.D., and A.J. Ridley. 2018. Rho GTPase signaling complexes in cell migration and invasion. J Cell Biol. 217:447–457.

44. Lorenz, M., H. Yamaguchi, Y. Wang, R.H. Singer, and J. Condeelis. 2004. Imaging sites of N-wasp activity in lamellipodia and invadopodia of carcinoma cells. Curr Biol. 14:697–703.

45. Machacek, M., L. Hodgson, C. Welch, H. Elliott, O. Pertz, P. Nalbant, A. Abell, G.L. Johnson, K.M. Hahn, and G. Danuser. 2009. Coordination of Rho GTPase activities during cell protrusion. Nature. 461:99–103.

46. Mahlandt, E.K., J.J.G. Arts, W.J. van der Meer, F.H. van der Linden, S. Tol, J.D. van Buul, T.W.J. Gadella, and J. Goedhart. 2021. Visualizing endogenous Rho activity with an improved localization-based, genetically encoded biosensor. J Cell Sci. 134.

47. Martin, K.H., K.E. Hayes, E.L. Walk, A.G. Ammer, S.M. Markwell, and S.A. Weed. 2012. Quantitative measurement of invadopodia-mediated extracellular matrix proteolysis in single and multicellular contexts. J Vis Exp:e4119.

48. Meirson, T., A. Genna, N. Lukic, T. Makhnii, J. Alter, V.P. Sharma, Y. Wang, A.O. Samson, J.S. Condeelis, and H. Gil-Henn. 2018. Targeting invadopodia-mediated breast cancer metastasis by using ABL kinase inhibitors. Oncotarget. 9:22158–22183.

49. Monteiro, P., C. Rossé, A. Castro-Castro, M. Irondelle, E. Lagoutte, P. Paul-Gilloteaux, C. Desnos, E. Formstecher, F. Darchen, D. Perrais, A. Gautreau, M. Hertzog, and P. Chavrier. 2013. Endosomal WASH and exocyst complexes control exocytosis of MT1-MMP at invadopodia. J Cell Biol. 203:1063–1079.

50. Moreau, V., F. Tatin, C. Varon, and E. Génot. 2003. Actin can reorganize into podosomes in aortic endothelial cells, a process controlled by Cdc42 and RhoA. Mol Cell Biol. 23:6809–6822.

51. Moshfegh, Y., J.J. Bravo-Cordero, V. Miskolci, J. Condeelis, and L. Hodgson. 2014. A Trio-Rac1-Pak1 signalling axis drives invadopodia disassembly. Nat Cell Biol. 16:574–586.

52. Murphy, D.A., and S.A. Courtneidge. 2011. The ’ins’ and ’outs’ of podosomes and invadopodia: characteristics, formation and function. Nat Rev Mol Cell Biol. 12:413–426.

53. Nagy, Z., K. Wynne, A. von Kriegsheim, S. Gambaryan, and A. Smolenski. 2015. Cyclic Nucleotide-dependent Protein Kinases Target ARHGAP17 and ARHGEF6 Complexes in Platelets. J Biol Chem. 290:29974–29983.

54. Nahidiazar, L., A.V. Agronskaia, J. Broertjes, B. van den Broek, and K. Jalink. 2016. Optimizing Imaging Conditions for Demanding Multi-Color Super Resolution Localization Microscopy. PLoS One. 11:e0158884.

55. Nakahara, H., S.C. Mueller, M. Nomizu, Y. Yamada, Y. Yeh, and W.T. Chen. 1998. Activation of beta1 integrin signaling stimulates tyrosine phosphorylation of p190RhoGAP and membrane-protrusive activities at invadopodia. J Biol Chem. 273:9–12.

56. Noll, B., D. Benz, Y. Frey, F. Meyer, M. Lauinger, S.A. Eisler, S. Schmid, P.L. Hordijk, and M.A. Olayioye. 2019. DLC3 suppresses MT1-MMP-dependent matrix degradation by controlling RhoB and actin remodeling at endosomal membranes. J Cell Sci.

57. Oikawa, T., T. Itoh, and T. Takenawa. 2008. Sequential signals toward podosome formation in NIH-src cells. J Cell Biol. 182:157–169.

58. Pan, S., Y. Deng, J. Fu, Y. Zhang, Z. Zhang, X. Ru, and X. Qin. 2018. Tumor Suppressive Role of ARHGAP17 in Colon Cancer Through Wnt/beta-Catenin Signaling. Cell Physiol Biochem. 46:2138–2148.

59. Pan, S.L., Y.Y. Deng, J. Fu, Y.H. Zhang, Z.J. Zhang, and X.J. Qin. 2022. ARHGAP17 enhances 5-Fluorouracil-induced apoptosis in colon cancer cells by suppressing Rac1. Neoplasma.

60. Pichot, C.S., C. Arvanitis, S.M. Hartig, S.A. Jensen, J. Bechill, S. Marzouk, J. Yu, J.A. Frost, and S.J. Corey. 2010. Cdc42-interacting protein 4 promotes breast cancer cell invasion and formation of invadopodia through activation of N-WASp. Cancer Res. 70:8347–8356.

61. Razidlo, G.L., B. Schroeder, J. Chen, D.D. Billadeau, and M.A. McNiven. 2014. Vav1 as a central regulator of invadopodia assembly. Curr Biol. 24:86–93.

62. Reinhard, N.R., M. Mastop, T. Yin, Y. Wu, E.K. Bosma, T.W.J. Gadella, Jr., J. Goedhart, and P.L. Hordijk. 2017. The balance between Gα(i)-Cdc42/Rac and Gα(1)(2)/(1)(3)-RhoA pathways determines endothelial barrier regulation by sphingosine-1-phosphate. Mol Biol Cell. 28:3371–3382.

63. Richnau, N., and P. Aspenstrom. 2001. Rich, a rho GTPase-activating protein domain-containing protein involved in signaling by Cdc42 and Rac1. J Biol Chem. 276:35060–35070.

64. Richnau, N., A. Fransson, K. Farsad, and P. Aspenström. 2004. RICH-1 has a BIN/Amphiphysin/Rvsp domain responsible for binding to membrane lipids and tubulation of liposomes. Biochem Biophys Res Commun. 320:1034–1042.

65. Rossman, K.L., C.J. Der, and J. Sondek. 2005. GEF means go: turning on RHO GTPases with guanine nucleotide-exchange factors. Nat Rev Mol Cell Biol. 6:167–180.

66. Sakurai-Yageta, M., C. Recchi, G. Le Dez, J.B. Sibarita, L. Daviet, J. Camonis, C. D’Souza-Schorey, and P. Chavrier. 2008. The interaction of IQGAP1 with the exocyst complex is required for tumor cell invasion downstream of Cdc42 and RhoA. J Cell Biol. 181:985–998.

67. Scott, R.W., D. Crighton, and M.F. Olson. 2011. Modeling and imaging 3-dimensional collective cell invasion. J Vis Exp.

68. Stoletov, K., and J.D. Lewis. 2015. Invadopodia: a new therapeutic target to block cancer metastasis. Expert Rev Anticancer Ther. 15:733–735.

69. Suman, P., S. Mishra, and H. Chander. 2018. High expression of FBP17 in invasive breast cancer cells promotes invadopodia formation. Med Oncol. 35:71.

70. Takano, K., K. Toyooka, and S. Suetsugu. 2008. EFC/F-BAR proteins and the N-WASP-WIP complex induce membrane curvature-dependent actin polymerization. Embo j. 27:2817–2828.

71. Tian, L., D.L. Nelson, and D.M. Stewart. 2000. Cdc42-interacting protein 4 mediates binding of the Wiskott-Aldrich syndrome protein to microtubules. J Biol Chem. 275:7854–7861.

72. Tian, Q., H. Gao, Y. Zhou, L. Zhu, J. Yang, B. Wang, P. Liu, and J. Yang. 2022. RICH1 inhibits breast cancer stem cell traits through activating kinases cascade of Hippo signaling by competing with Merlin for binding to Amot-p80. Cell Death & Disease. 13:71.

73. Timmerman, I., N. Heemskerk, J. Kroon, A. Schaefer, J. van Rijssel, M. Hoogenboezem, J. van Unen, J. Goedhart, T.W. Gadella, Jr., T. Yin, Y. Wu, S. Huveneers, and J.D. van Buul. 2015. A local VE-cadherin and Trio-based signaling complex stabilizes endothelial junctions through Rac1. J Cell Sci. 128:3041–3054.

74. Tonucci, F.M., E. Almada, C. Borini-Etichetti, A. Pariani, F. Hidalgo, M.J. Rico, J. Girardini, C. Favre, J.R. Goldenring, M. Menacho-Marquez, and M.C. Larocca. 2019. Identification of a CIP4 PKA phosphorylation site involved in the regulation of cancer cell invasiveness and metastasis. Cancer Lett.

75. Tsujita, K., A. Kondo, S. Kurisu, J. Hasegawa, T. Itoh, and T. Takenawa. 2013. Antagonistic regulation of F-BAR protein assemblies controls actin polymerization during podosome formation. J Cell Sci. 126:2267–2278.

76. Tsujita, K., S. Suetsugu, N. Sasaki, M. Furutani, T. Oikawa, and T. Takenawa. 2006. Coordination between the actin cytoskeleton and membrane deformation by a novel membrane tubulation domain of PCH proteins is involved in endocytosis. J Cell Biol. 172:269–279.

77. van Unen, J., T. Yin, Y.I. Wu, M. Mastop, T.W. Gadella, Jr., and J. Goedhart. 2016. Kinetics of recruitment and allosteric activation of ARHGEF25 isoforms by the heterotrimeric G-protein Gαq. Sci Rep. 6:36825.

78. Vaughan, E.M., A.L. Miller, H.Y. Yu, and W.M. Bement. 2011. Control of local Rho GTPase crosstalk by Abr. Curr Biol. 21:270–277.

79. Vinci, M., C. Box, and S.A. Eccles. 2015. Three-dimensional (3D) tumor spheroid invasion assay. J Vis Exp:e52686.

80. Wells, C.D., J.P. Fawcett, A. Traweger, Y. Yamanaka, M. Goudreault, K. Elder, S. Kulkarni, G. Gish, C. Virag, C. Lim, K. Colwill, A. Starostine, P. Metalnikov, and T. Pawson. 2006. A Rich1/Amot complex regulates the Cdc42 GTPase and apical-polarity proteins in epithelial cells. Cell. 125:535–548.

81. Yamaguchi, H. 2012. Pathological roles of invadopodia in cancer invasion and metastasis. Eur J Cell Biol. 91:902–907.

82. Yamaguchi, H., M. Lorenz, S. Kempiak, C. Sarmiento, S. Coniglio, M. Symons, J. Segall, R. Eddy, H. Miki, T. Takenawa, and J. Condeelis. 2005. Molecular mechanisms of invadopodium formation: the role of the N-WASP-Arp2/3 complex pathway and cofilin. J Cell Biol. 168:441–452.

83. Yamamoto, H., M. Sutoh, S. Hatakeyama, Y. Hashimoto, T. Yoneyama, T. Koie, H. Saitoh, K. Yamaya, T. Funyu, T. Nakamura, C. Ohyama, and S. Tsuboi. 2011. Requirement for FBP17 in invadopodia formation by invasive bladder tumor cells. J Urol. 185:1930–1938.

84. Yu, X., T. Zech, L. McDonald, E.G. Gonzalez, A. Li, I. Macpherson, J.P. Schwarz, H. Spence, K. Futó, P. Timpson, C. Nixon, Y. Ma, I.M. Anton, B. Visegrády, R.H. Insall, K. Oien, K. Blyth, J.C. Norman, and L.M. Machesky. 2012. N-WASP coordinates the delivery and F-actin-mediated capture of MT1-MMP at invasive pseudopods. J Cell Biol. 199:527–544.

85. Zagryazhskaya-Masson, A., P. Monteiro, A.S. Macé, A. Castagnino, R. Ferrari, E. Infante, A. Duperray-Susini, F. Dingli, A. Lanyi, D. Loew, E. Génot, and P. Chavrier. 2020. Intersection of TKS5 and FGD1/CDC42 signaling cascades directs the formation of invadopodia. J Cell Biol. 219.

